# Naturally occurring mutations of SARS-CoV-2 main protease confer drug resistance to nirmatrelvir

**DOI:** 10.1101/2022.06.28.497978

**Authors:** Yanmei Hu, Eric M. Lewandowski, Haozhou Tan, Xiaoming Zhang, Ryan T. Morgan, Xiujun Zhang, Lian M. C. Jacobs, Shane G. Butler, Maura V. Gongora, John Choy, Xufang Deng, Yu Chen, Jun Wang

## Abstract

The SARS-CoV-2 main protease (M^pro^) is the drug target of Pfizer’s oral drug Paxlovid. The emergence of SARS-CoV-2 variants with mutations in M^pro^ raised the alarm of potential drug resistance. In this study, we identified 100 naturally occurring M^pro^ mutations located at the nirmatrelvir binding site, among which 20 mutants, including S144M/F/A/G/Y, M165T, E166G, H172Q/F, and Q192T/S/L/A/I/P/H/V/W/C/F, showed comparable enzymatic activity to the wild-type (k_cat_/K_m_ <10-fold change) and resistance to nirmatrelvir (K_i_ >10-fold increase). X-ray crystal structures were determined for seven representative mutants with and/or without GC-376/nirmatrelvir. Viral growth assay showed that M^pro^ mutants with reduced enzymatic activity led to attenuated viral replication. Overall, our study identified several drug resistant hot spots that warrant close monitoring for possible clinical evidence of Paxlovid resistance.

**One Sentence Summary:** Paxlovid resistant SARS-CoV-2 viruses with mutations in the main protease have been identified from clinical isolates.

## INTRODUCTION

The ongoing COVID-19 pandemic highlights the urgent need of orally bioavailable antiviral drugs. Paxlovid is a combination of the viral main protease (M^pro^ or 3CL^pro^) inhibitor nirmatrelvir and the metabolic booster ritonavir (*1, 2*). M^pro^ is a cysteine protease that mediates the cleavage of viral polyproteins during viral replication and is a high-profile antiviral drug target (*3*). In addition to Paxlovid, other M^pro^ inhibitors advanced to the clinical stage include PF-07304814 (phosphate form of PF-00835231), S-217622, PBI-0451, EDP-235, and 13b (*4*). As an RNA virus, SARS-CoV-2 continues to evolve with or without selection pressure. The recent emergence of variants of concern, particularly the Omicron variant, raises the concern of possible altered susceptibility to vaccines and antiviral drugs. In this study, we report the discovery of drug resistant M^pro^ mutants from naturally occurring SARS-CoV-2 polymorphisms deposited in the Global Initiative on Sharing Avian Influenza Data (GISAID) database. Three inhibitors - GC-376, PF-00835231, and nirmatrelvir - were examined for drug resistance. GC-376 is a veterinary drug candidate for the treatment of feline infectious peritonitis virus (FIPV) infection in cats (*5*). The phosphonate prodrug of PF-00835231 was a clinical candidate developed by Pfizer as an intravenous drug for the treatment of COVID patients in hospitals (*6*). Predicting drug resistance before it becomes dominant in clinic is vital in facilitating antiviral drug development to combat the pandemic.

## RESULTS

### Identification of SARS-CoV-2 M^pro^ mutants from GISAID sequence analysis

Recent sequence analysis of SARS-CoV-2 M^pro^ revealed multiple prevalent mutations including G15S, T21I, L89F, K90R, P108S, P132H, and L205V (*7-9*). All these mutants are located outside the nirmatrelvir binding site (Fig. 1A) and were found to have similar catalytic efficacy (k_cat_/K_m_) as the wild-type (WT) protein (*7, 10*). These mutants remained susceptible to nirmatrelvir with no significant IC_50_ or K_i_ value shifts (< 2-fold) (*7*). Nevertheless, drug resistance to nirmatrelvir is anticipated given the experience from the clinical use of HIV and HCV protease inhibitors (*11, 12*). Several studies have been conducted to evolve or predict nirmatrelvir resistant M^pro^ mutants in cell culture (*13-16*). In this study, we took a different approach by discovering drug resistant M^pro^ mutants from circulating SARS-CoV-2 variants.

**Fig. 1.**
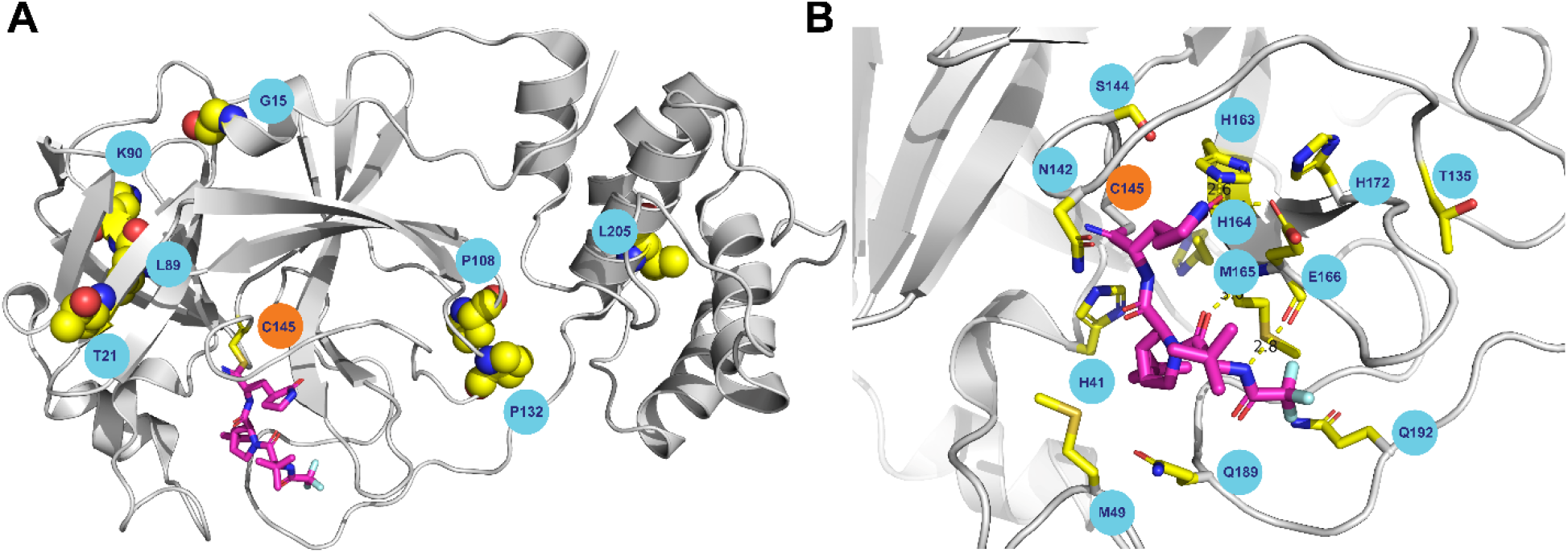
SARS-CoV-2 M^pro^ mutants identified from GISAID sequence analysis. (**A**) Residues with high mutation rates that were previously examined. None of the mutants showed significant drug resistance. (**B**) Residues located within 6 Å of the nirmatrelvir binding site that were examined in this study. Figures were generated using Pymol with the X-ray crystal structure of nirmatrelvir in complex with SARS-CoV-2 M^pro^ (PDB: 7SI9). Nirmatrelvir is colored in magenta.

To identify drug resistant mutants of M^pro^, we focus on the active site residues that are located within 6 Å of the nirmatrelvir binding site (PDB: 7SI9) (*2*). In total, 12 residues were selected including H41, M49, T135, N142, S144, H163, H164, M165, E166, H172, Q189, and Q192 (Fig. 1B). We expect that mutations at these active site residues will have a direct impact on substrate binding and drug inhibition. To test this hypothesis, we analyzed the mutations of these 12 residues using the SARS-CoV-2 sequences deposited in GISAID (*17*) and the mutation frequency of each active site residue was plotted in Fig. 2A.

**Fig. 2.**
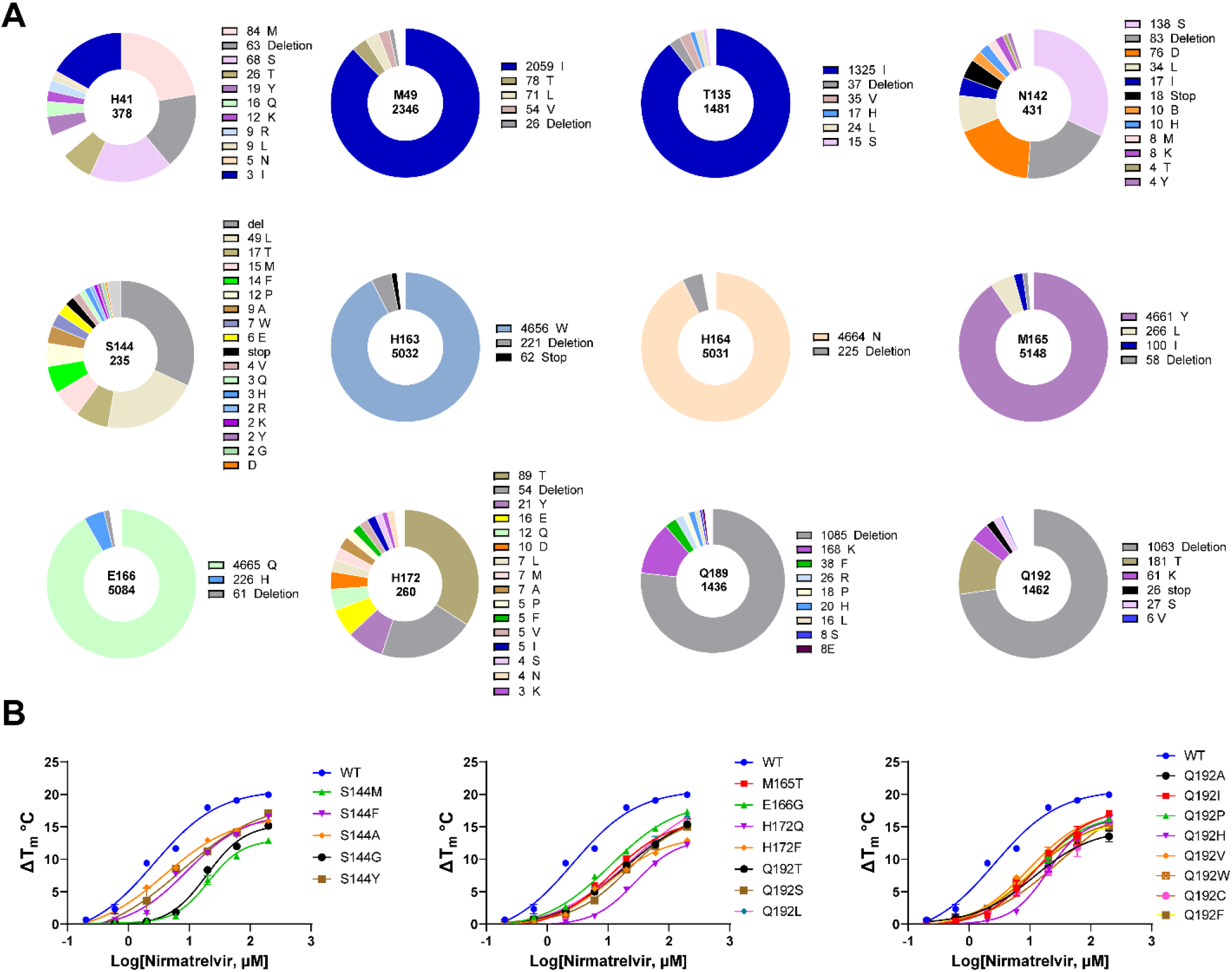
SARS-CoV-2 M^pro^ mutants characterized in this study. (**A**) Mutations at 12 residues located at the nirmatrelvir binding site. Sequence data were obtained from CoVsurver of the GISAD, developed by A*STAR Bioinformatics Institute (BII), Singapore. A total of 5,124,505 mutations of Nsp5 (M^pro^) were obtained from the database as of June 21, 2022. Occurrence of total mutations for each amino acid residue is labeled in the center of each pie chart. Occurrences of specific mutations are labeled on the right of its affiliate pie chart. (**B**) Characterization of nirmatrelvir resistance against M^pro^ mutants by the thermal shift assay. The results are the average of duplicates.

Based on the sequence analysis, we chose 100 mutants that cover all high frequency mutations at these 12 residues. The tag-free recombinant SARS-CoV-2 M^pro^ mutants with native N- and C-termini were expressed in *E. Coli* and purified (fig. S1). All proteins were folded correctly as shown by the thermal shift assay (fig. S2) and the enzymatic assay (table S1). The enzymatic activity (k_cat_/K_m_) of the mutant proteins was determined using the FRET assay (*18, 19*). To profile the drug resistance, we tested the M^pro^ mutants against nirmatrelvir in the FRET assay and determined the half maximal inhibitory concentration (IC_50_) and inhibition constant (K_i_). For mutants showing resistance against nirmatrelvir, the drug sensitivity was further tested against PF-00835231 and GC-376 for cross-resistance. The comprehensive data set is shown in table S1.

### H41 and H163 are critical for the enzymatic activity

Among the 100 mutants, H41M, H41T, H41Y, and H163W were enzymatically inactive (table S1, yellow). H41 forms the catalytic dyad with C145, and all three mutants – H41M (84 occurrences), H41T (27 occurrences), and H41Y (19 occurrences) – were detrimental to the enzymatic activity (table S1). The X-ray crystal structure showed that the side chain imidazole of H163 forms a hydrogen bond with the carbonyl from the nirmatrelvir P1 pyrrolidone (Fig. 1B) (PDB: 7SI9) (*2*) or similar functional groups from other inhibitors, suggesting its essential contribution to drug binding. Although H163W is a high frequency mutation with 4,672 occurrences, this mutant led to inactive enzyme.

### S144, M165, E166, H172, and Q192 are hot spots for drug resistance

It is generally assumed that mutations with impaired enzymatic activity will lead to attenuation of viral replication. We therefore focused on M^pro^ mutants with k_cat_/K_m_ values within 10-fold variation compared to WT. For drug sensitivity, a K_i_ increase by more than 10-fold is defined as significant resistance. In total, 20 M^pro^ mutants met both criteria including S144M, S144F, S144A, S144G, S144Y, M165T, E166G, H172Q, H172F, Q192T, Q192S, Q192L, Q192A, Q192I, Q192P, Q192H, Q192V, Q192W, Q192C, and Q192F (table S1, red).

S144 locates at the S1 pocket and is part of the oxyanion hole consisting of two additional residues G143 and C145 (Fig. 1B). Among the top 15 high frequency mutations, five mutants - S144M (8.0-fold lower k_cat_/K_m_), S144F (5.8-fold), S144A (1.8-fold), S144G (2.6-fold), and S144Y (7.8-fold) - had comparable enzymatic activity to the WT. Significantly, all five mutants showed drug resistance against nirmatrelvir with K_i_ values increased between 19.2 to 38.0-fold. Pfizer’s report for healthcare providers similarly disclosed S144A as a nirmatrelvir resistant mutant with a K_i_ increase of 91.9-fold (*20*) compared to 20.5-fold from our study. Four mutants - S144L (183.3-fold lower in k_cat_/K_m_), S144P (523.8-fold), S144R (478.3-fold), and S144K (534.0-fold) - had significantly reduced enzymatic activity compared to WT and increased resistance to nirmatrelvir. Similarly, the remaining seven mutants - S144T, S144W, S144E, S144V, S144Q, S144H, and S144D - had compromised enzymatic activity with k_cat_/K_m_ values decreased between 20.0 to 85.9-fold compared to WT. A significant drug resistance against nirmatrelvir was also observed for these seven mutants.

M165 locates at the S2 pocket and forms a hydrophobic interaction with the P2 dimethylcyclopropylproline (Fig. 1B). The most frequent mutant, M165Y, had a significantly reduced enzymatic activity (41.7-fold decrease in k_cat_/K_m_), while M165L, M165I, M165V, M165T, M165A, and M165C had similar enzymatic activity as the WT. No drug resistance was observed for M165L, M165I, M165V, M165A, and M165C. However, a significant drug resistance against nirmatrelvir was observed for M165T (29.9-fold increase in K_i_). The remaining mutants M165W/K/R/G/F/H/P/D had a significantly reduced enzymatic activity (>14-fold decrease in k_cat_/K_m_).

E166 locates at the S1 pocket and forms three hydrogen bonds with nirmatrelvir (Fig. 1B). E166Q is a high frequency mutation with 4,681 occurrences. It has a similar enzymatic activity (k_cat_/K_m_) as the WT, and no significant drug resistance against nirmatrelvir was observed (4.5-fold increase in K_i_). E166H/K/L/Y/I mutants all had a significant reduction of enzymatic activity (>17.5-fold decrease in k_cat_/K_m_) and a high degree of drug resistance. Interestingly, E166G only had a 7.4-fold reduction in enzymatic activity k_cat_/K_m_ value but a 16.4-fold increase in K_i_ value against nirmatrelvir. In parallel to our study, two preprints reported nirmatrelvir resistant M^pro^ mutants identified from serial viral passage experiments in cell culture (*13, 14*). Jochmans et al. discovered a triple mutant, L50F/E166A/L167F, with a 72-fold IC_50_ increase in the enzymatic assay and a 51-fold EC_50_ increase in the antiviral assay against nirmatrelvir (*14*). However, the triple mutant only had 5.3% of the enzymatic activity of the WT, indicating an impaired fitness of replication. In another study, Zhou et al. showed that the L50F/E166V double mutant led to an 80-fold resistance in the antiviral assay against nirmatrelvir (*13*). Both E166A and E166V are naturally occurring mutations with 5 and 7 occurrences. Taken together, E166 appears to be a hotspot for drug resistant mutation.

H172 locates at the S1 pocket but does not directly interact with nirmatrelvir (Fig. 1B). Among the 17 H172 mutants examined, H172Q (3.2-fold lower k_cat_/K_m_) and H172F (9.9-fold) had comparable enzymatic activity as the WT. Both mutants also showed a significant drug resistance against nirmatrelvir (>42-fold increase in K_i_). The H172Y (13.9-fold lower k_cat_/K_m_) and H172A (11.3-fold lower k_cat_/K_m_) mutants had reduced enzymatic activity, while being resistant to nirmatrelvir (>113.7-fold increase in K_i_). H172Y was similarly disclosed by Pfizer as a nirmatrelvir resistant mutant (233-fold increase in K_i_) (*20*). The remaining mutants H172T/E/D/L/M/I/V/S/N/K/R/G/C had a significantly reduced enzymatic activity (>21.0-fold lower k_cat_/K_m_).

Q192 locates at the S4 pocket and forms a hydrophobic interaction with the trifluoromethyl substitution from nirmatrelvir (Fig. 1B). Q192T (9.2-fold lower k_cat_/K_m_), Q192S (8.9-fold), Q192L(4.3-fold), Q192A (6.2-fold), Q192I (5.6-fold), Q192P (7.6-fold), Q192H (8.2-fold), Q192V (9.0-fold), Q192W (8.0-fold), Q192C (7.0-fold), and Q192F (3.5-fold) had comparable enzymatic activity as the WT, and all showed resistance against nirmatrelvir (>22.2-fold increase in K_i_). Cross resistance was also observed with PF-00835231 (>25.5-fold) and GC-376 (>7.7-fold).

The drug resistance of these 20 M^pro^ mutants against nirmatrelvir was further confirmed in the thermal shift drug titration assay. All mutants displayed a lower degree of protein stabilization than the WT M^pro^ with increasing concentrations of nirmatrelvir (Fig. 2B).

### M49, T135, N142, H164, M165, and Q189 can tolerate multiple mutations without significantly affecting enzymatic activity and drug inhibition

The most frequent mutants at residue M49 - M49I, M49T, M49L, and M49V - remained sensitive to nirmatrelvir (< 3-fold change in IC_50_). Interestingly, the enzymatic activity (k_cat_/K_m_) of the M49I and M49L mutants showed 1.69 and 1.74-fold increase compared to WT.

T135I is a high frequency mutation with 1,340 occurrences. The T135I mutant had a similar k_cat_/K_m_ value as the WT and remained sensitive to all three inhibitors (<2.9-fold change in K_i_).

The top nine high frequency mutants at residue N142 all had similar enzymatic activity as the WT (<4.1-fold change in k_cat_/K_m_) and remained sensitive to nirmatrelvir (<3.5-fold change in IC_50_).

H164N is a high frequency mutation with 4,681 occurrences and remained sensitive to all three inhibitors (<4.1-fold change in K_i_). The catalytic activity of the H164N mutant (4.2-fold lower in k_cat_/K_m_) is comparable to WT.

All eight Q189 mutants retained similar enzymatic activity as the WT with the change in k_cat_/K_m_ values between 1.9- and 9.2-fold. No significant drug resistance was observed for nirmatrelvir (<3.1-fold change in IC_50_). Interestingly, the enzymatic activity of Q189E increased by nearly 2-fold compared to WT.

Collectively, the results suggest that M49, T135, N142, H164, and Q189 might be able to accommodate multiple mutants without a significant compromise in enzymatic activity and drug sensitivity.

### H172Y/Q189E double mutant rescued the enzymatic activity and maintained drug resistance

Given the enhanced enzymatic activity of the Q189E mutant (1.9-fold increase in k_cat_/K_m_) and the reduced enzymatic activity of the H172Y mutant (13.9-fold decrease) compared to WT (fig. 4A), we hypothesized that the H172Y/Q189E double mutant might rescue the reduced enzymatic activity of the drug resistant H172Y mutant. Indeed, the H172Y/Q189E double mutant increased both the enzymatic activity of H172Y from 790 M^-1^S^-1^ to 1,009 M^-1^S^-1^ (fig. 4A) and resistance against nirmatrelvir (281.1-fold increase in K_i_ value in comparison to 146.3-fold increase of H172Y single mutation) (fig. 4B). The resistance of the H172Y/Q189E against nirmatrelvir was further confirmed in the thermal shift binding assay (fig. 4C). These results suggest that although many of the identified single mutants had reduced enzymatic activity (table S1), it is plausible that the virus can generate multiple mutations to help restore the fitness of replication while maintaining, or even enhancing drug resistance.

### Recombinant SARS-CoV-2 viruses with M^pro^ S144A and H172Y mutants had attenuated replication in cell culture

To investigate the effect of M^pro^ mutants on viral replication and the sensitivity to nirmatrelvir, we chose two representative mutants S144A and H172Y. To this end, we successfully generated two recombinant SARS-CoV-2 viruses harboring the S144A and H172Y mutations, designated rNsp5^S144A^ and rNsp5^H172Y^, respectively, using a SARS-CoV-2 reverse genetic system (*21*). An isogenic wild-type recombinant SARS-CoV-2 WA1 strain (rSARS2-WT) was also generated and served as a wild-type control. The nsp5 coding sequences of these recombinant viruses were sequenced and the corresponding mutants were confirmed. We performed plaque assay and growth kinetics analysis to evaluate viral replication. As shown in Fig. 3A, rSARS-CoV-2 Nsp5^S144A^ mutant formed slightly smaller plaques than the rSARS2-WT, whereas the plaques of rNsp5^H172Y^ mutant were drastically smaller than those of rSARS2-WT and rNsp5^S144A^. Growth kinetics analysis revealed that the rNsp5^S144A^ produced a 1-log lower extracellular virus titer at the exponential growth phase (Fig. 3B). In stark contrast, the rNsp5^H172Y^ mutant exhibited significant replication defect and had 2∼3-log lower titers at the exponential growth phase compared to rSARS2-WT. These data together demonstrate that the S144A mutation mildly impaired viral growth, whereas the H172Y mutation severely weakens viral replication, consistent with the *in vitro* enzymatic analysis results (table S1). To assess the sensitivity to nirmatrelvir, we performed antiviral experiments using a Celltiter Glo assay to measure the virus-induced cytopathic effect in Vero-E6 cells expressing hACE2 and hTMPRSS2. As shown in Fig. 3C, the EC_50_ value of rNsp5^S144A^ was 366.9 nM, 2.4-fold higher than the EC_50_ (152.6 nM) of rSARS2-WT. The antiviral assay for the rNsp5^H172Y^ mutant was not successful due to its severe replication defect. Taken together, the results from these live virus analyses are consistent with our enzymatic assay result: M^pro^ mutants with a significantly reduced enzymatic activity such as H172Y (13.9-fold decrease in k_cat_/K_m_) had a high fitness cost, while mutants with comparable enzymatic activity as WT such as S144A (1.8-fold decrease in k_cat_/K_m_) had similar growth kinetics as the WT.

**Fig. 3.**
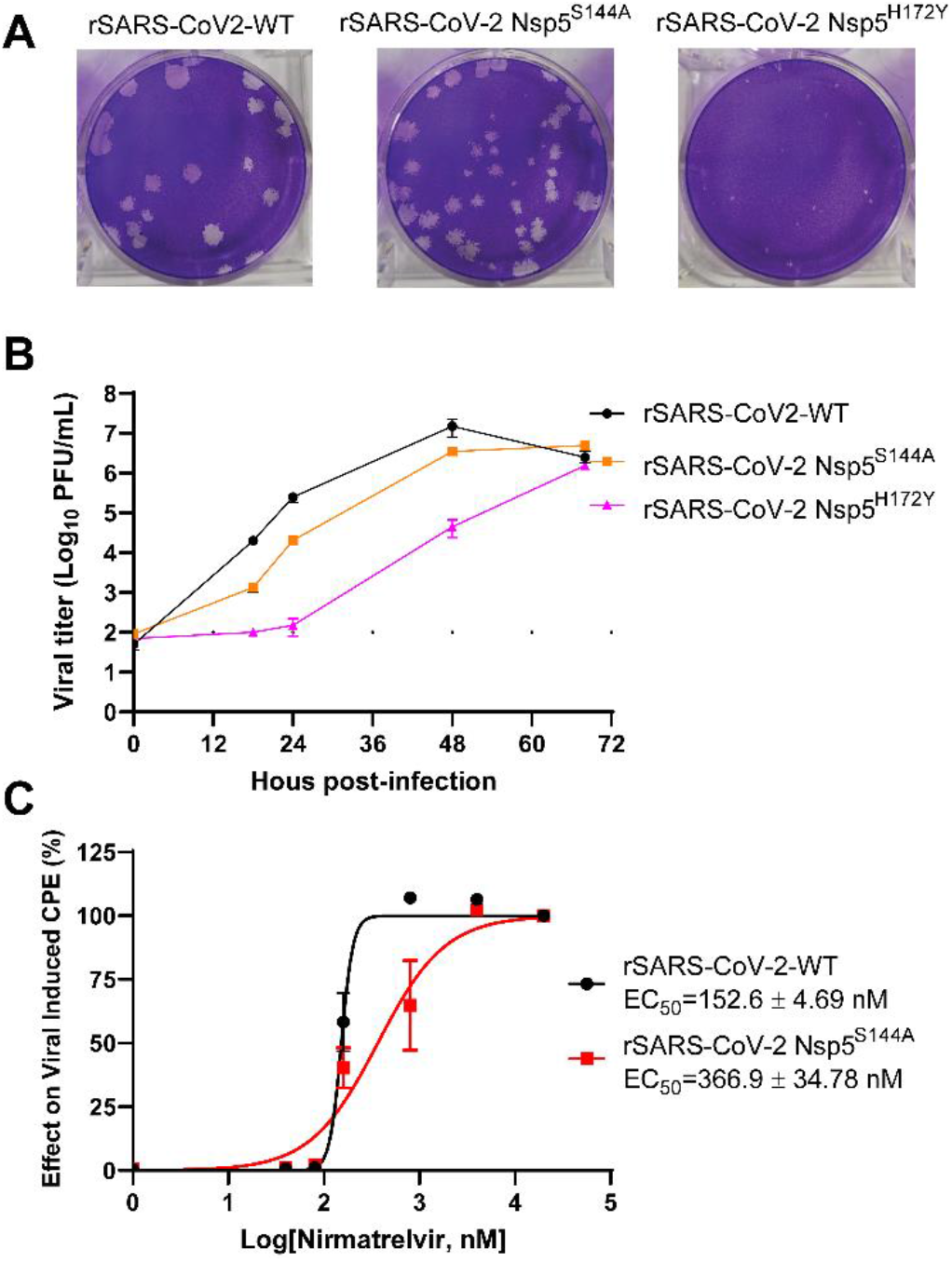
Characterization of the recombinant SARS-CoV-2 Nsp5^S144A^ and Nsp5^H172Y^ viruses. (**A**) Plaque formation by recombinant SARS-CoV-2 wild-type (rSARS-CoV-2-WT) and the M^pro^ mutant virus rNsp5-S144A and rNsp5-H172Y. The H172Y mutant virus formed significantly smaller plaques. (**B**) Growth kinetics of recombinant WT virus and the Nsp5 mutant viruses in Vero-E6 expressing hACE2 and hTMPRSS2. (**C**) Antiviral assay in Vero-E6 expressing hACE2 and hTMPRSS2. The dotted line represents the limit of detection. All experiments were independently performed at least twice. Data are shown as mean ± SD.

**Fig. 4.**
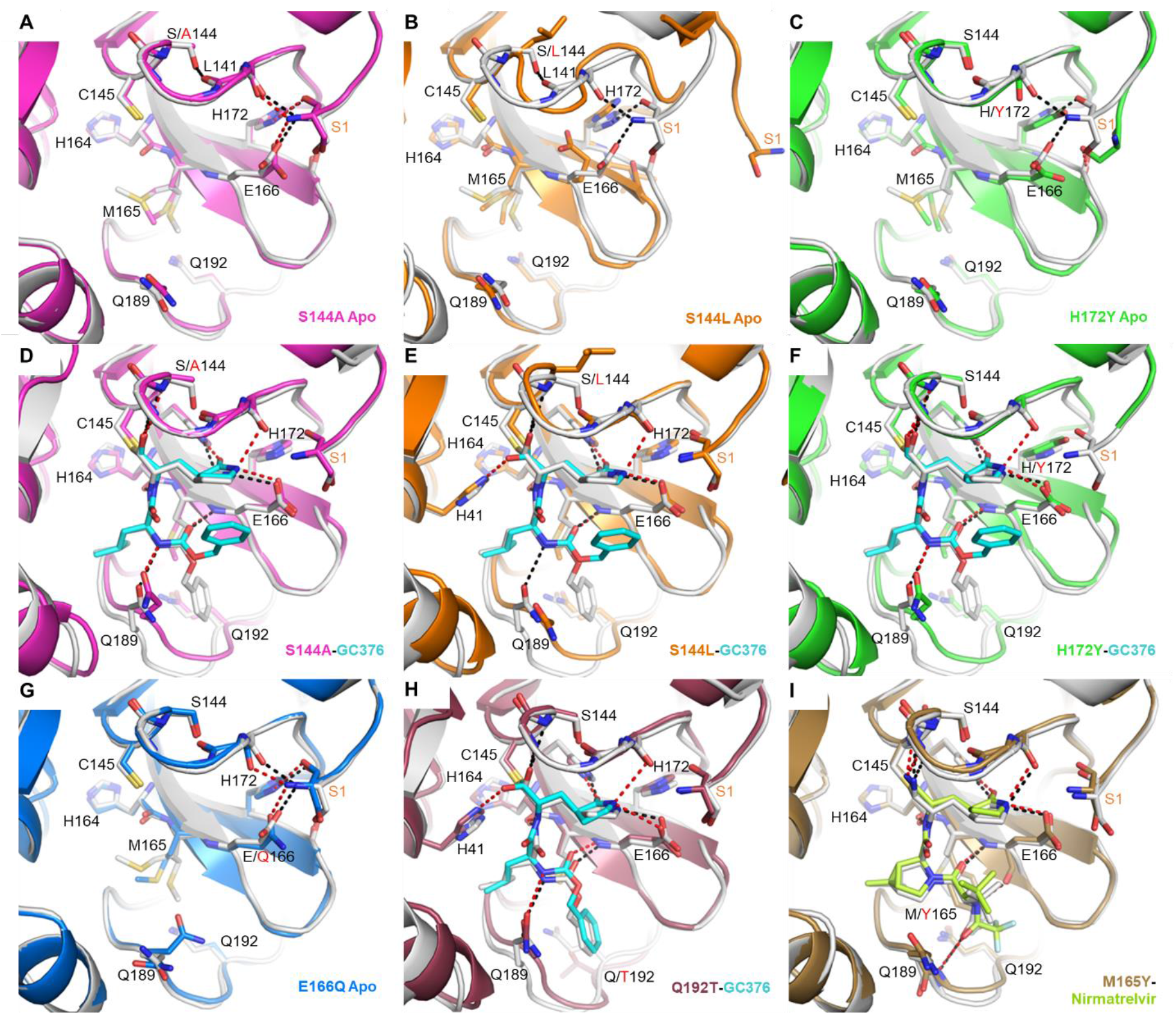
X-ray crystal structures of SARS-CoV-2 M^pro^ mutants. Each mutant structure is aligned with the corresponding WT structure shown in white (apo, PDB 7JP1; GC-376 complex, PDB 6WTT; nirmatrelvir complex, PDB 7RFW). For the mutant structures, GC-376 and nirmatrelvir are shown in cyan and neon green respectively. WT HBs are shown as black dashes for selected residues at the mutation sites or between the protein and inhibitor. Mutant HBs are shown as red dashes. Mutations are indicated with red text. The S1 residue from the N-terminus of the adjacent protomer is labeled in orange. The side chain of L141 is not shown. (**A**) Apo M^pro^ S144A (magenta, PDB 8D4L). (**B**) Apo M^pro^ S144L (orange, PDB 8DFE). The view for panel B is shifted slightly to show the movement of the adjacent N-terminus. (**C**) Apo Mpro H172Y (green, PDB 8D4J). (**D**) M^pro^ S144A GC-376 complex (magenta, PDB 8D4M). (**E**) M^pro^ S144L GC-376 complex (orange, PDB 8DD9). (**F**) M^pro^ H172Y GC-376 complex (green, PDB 8D4K). S1 residue is disordered and not modeled. (**G**) Apo M^pro^ E166Q (blue, PDB 8D4N). (**H**) M^pro^ Q192T GC376 complex (mauve, PDB 8DGB). (**I**) M^pro^ M165Y nirmatrelvir complex (brown, PDB 8DCZ).

### Structural basis for resistance mutations

We have determined the X-ray crystal structures of several key representative mutants, including both unbound and GC-376 complex structures for S144A, S144L, H172Y, the GC-376 complex of Q192T, and the nirmatrelvir complex of M165Y, at 1.70-2.87 Å resolutions (Fig. 4, table S2). The structures of H164N (apo and GC-376 bound) and E166Q (apo) were also determined for comparison (fig. S5). GC-376 and nirmatrelvir place the same pyrrolidone group in the S1 pocket. It should be noted that the terminal benzene group of GC-376 exhibited two different binding modes in previously published WT structures (*18, 19*). The conformational difference in this substitution between mutants (except Q192T) and the WT likely originated from its flexibility, rather than the specific mutations.

All the WT residues at the four mutation sites (S144, M165, H172, and Q192) are involved in intra-molecular interactions and are at least partially buried. However, except for S144L, the changes caused by the five mutations are mostly local and small. All mutant complex structures including S144L showed the inhibitor and the protein assumed conformations very similar to the WT.

The decrease in the mutants’ catalytic activity and inhibition by GC-376/nirmatrelvir appears to stem from two causes, a largely enthalpic effect through direct disruption of ligand-binding interactions and an entropic effect through increasing conformational instability of the active site. S144L, M165Y, H172Y and Q192T represented some of the biggest changes in both the residue size and the enzyme activity, with a decrease of ∼127x, 39x, 13x, and 10x respectively in k_cat_ values. Those mutations resulted in notable changes in ligand interactions in the structures. In the unbound state (Fig. 5B) and, to a lesser extent, the complex structure (Fig. 4E), the S144L mutation led to a drastically different conformation in the 140-146 loop, as well as neighboring regions to avoid steric clashes caused by the bulky leucine side chain. Importantly, this loop constitutes the oxyanion hole stabilizing the transition state of the enzymatic reaction, formed by the main chain amide groups of G143, S144, and C145. Interestingly, in the GC-376 complex structure of S144L (Fig. 4E) as well as Q192T (Fig. 4H), the thioacetal hydroxide is placed outside the oxyanion hole and hydrogen bonds with H41, unlike most previously determined structures (*18*). This suggests that the interactions between the oxyanion hole and the inhibitor thioacetal hydroxide, as well as, by extension, the substrate transition state, are likely weakened in the S144L and Q192T mutants, resulting in an alternate conformation in the inhibitor crystal structure. Similarly, the M165Y mutation also seemed to lead to diminished interactions between the oxyanion hole and the ligand, indicated by the lengthened HB between the imine nitrogen and G143 amide NH (from 3.0 Å in the WT to 3.6 Å in the M165Y mutant) (Fig. 4I). These distortions are likely caused by the bulky Y165 residue through series of ripple effects relayed by both the protein and inhibitor in a tightly packed active site. Meanwhile, the H172Y mutation altered the conformation of the N-terminus of the adjacent protomer in the biological dimer, which is near the active site. In addition to these effects on the reaction center, the M165Y and Q192T mutations also directly impacted ligand interactions in the S4 site. The M165Y mutation pushes the terminal trifluoromethyl group of nirmatrelvir out of its original position in the WT complex, disrupting its interaction with L167 and the T190 backbone oxygen (Fig. 4I). The Q192T mutation increased the plasticity of the surrounding residues (Fig. 4H), allowing them to better accommodate the terminal benzene ring of GC-376, which assumed a different conformation compared with the other mutants.

In contrast to the above four mutations, the remaining resistant mutation, S144A, represented the smallest changes in the side chain size, and led to nearly no alteration of the unbound structure compared with the WT, similar to the structure of the E166Q mutant that remained sensitive to nirmatrelvir inhibition (Figs. 4A, D, G). However, the S144 mutant located in the active site, abolishes the intramolecular interactions involving S144 side chain in the WT, and subsequently increase the conformational instability of the protein active site. Even though the enthalpic interactions between the substrate/inhibitor and the protein may be similar to the WT in the lowest energy conformation, the entropic cost will be higher for the mutants, thus making the free energy of ligand binding less favorable. As the inhibitor relies on better shape complementarity with a smaller portion of the active site and contains more rigid features than the substrate, we hypothesize that this entropic cost may impact inhibitor binding more than the larger and more flexible substrate. For example, compared with the glutamine side chain at the P1 position, the pyrrolidone ring of nirmatrelvir and GC-376 forms additional interactions with the peptide bond between L141 and N142 through the two extra carbon atoms. The S144A mutation eliminates the HB with the carbonyl group of this peptide bond, likely increasing its flexibility and increasing the energetic cost of inhibitor binding. Such entropic effects are not limited to the S144A mutation causing minimal structural changes, but also apply to those aforementioned mutations such as S144L and H165Y that can both directly influence the protein-ligand contacts and increase the active site flexibility.

## DISCUSSION

Collectively, our results have several implications. First, all 100 M^pro^ mutants characterized in this study are naturally occurring SARS-CoV-2 M^pro^ polymorphisms that could potentially affect the efficacy of Paxlovid, and continuous prescription of Paxlovid might likely increase the frequency of these pre-existing drug resistance mutants. Second, S144, M165, E166, H172, and Q192 appear to be hot spots for nirmatrelvir resistance and need to be closely monitored among circulating viruses. Mutations at these residues are most likely to maintain the enzymatic activity while causing a significant drug resistance. Third, M^pro^ mutants with a significantly reduced enzymatic activity (>10-fold decrease in k_cat_/K_m_) such as H172Y impairs the fitness of viral replication in cell culture, suggesting the short-term resistance risk due to single mutations may be relatively low. Fourth, as exemplified by the H172Y/Q189T mutant, it is possible that double or triple mutants might emerge to compensate for the loss of enzymatic activity from the single mutant while maintaining or enhancing drug resistance, which significantly raise the near and long-term resistance risk especially considering the multiple naturally occurring mutations shown by our study to confer resistance. Therefore, the M^pro^ mutants with reduced enzymatic activity from this study should also be monitored.

## MATERIALS AND METHDOS

### Materials

Oligonucleotides were from Integrated DNA Technologies (Coralville, IA). The SARS-CoV-2 M^pro^ FRET substrate Dabcyl-KTSAVLQ/SGFRKME-(Edans) was synthesized as previously described (*18*). All other chemicals were purchased from either Sigma Aldrich (St. Louis, MO) or Fisher Scientific (Pittsburgh, PA). DNA sequencing was performed by Azenta Life Sciences (South Plainfield, NJ). The following reagent was obtained through BEI Resources, NIAID, NIH: Cercopithecus aethiops Kidney Epithelial Cells Expressing Transmembrane Protease, Serine 2 and Human Angiotensin-Converting Enzyme 2 (Vero E6-TMPRSS2-T2A-ACE2), NR-54970; SARS-Related Coronavirus 2, USA-WA1/2020 Recombinant Infectious Molecular Clone Plasmid Kit, NR-53762. Vero E6-TMPRSS2-T2A-ACE2 cells were maintained in Dulbecco’s modified Eagle medium (DMEM) (Corning, 10013CM) containing 10% heat-inactivated fetal bovine serum (FBS), 1% Pen/Strep, 1× nonessential amino acid, and 10 μg/mL puromycin (InVivogen, ant-pr-1) to maintain the expression of TMPRSS2 and ACE2. The SARS-CoV-2 infectious plasmid clones were propagated in bacterial cells TOP10 strain (ThermoFisher, C404010) and sequenced.

### SARS-CoV-2 M^pro^ Mutagenesis, Protein Expression and Purification

SARS-CoV-2 M^pro^ mutants were generated with QuikChange® II Site-Directed Mutagenesis Kit from [Agilent (Catalog #200524](https://community.agilent.com/?banner=web.acom-footer.community)), using previously created plasmid pE-SUMO-M^pro^ (from strain BetaCoV/Wuhan/WIV04/2019) (*19*) as the template. The plasmid produces tag-free M^pro^ protein with no extra residue at either N or C terminus upon removal of the SUMO tag by SUMO protease digestion.

SARS-CoV-2 M^pro^ mutant proteins were expressed and purified as previously described (*18, 19*) with minor modifications. Plasmids were transformed into *E. coli* BL21(DE3) competent cells and bacterial cultures overexpressing the target proteins were grown in LB (Luria-Bertani) medium containing 50 µg/mL of kanamycin at 37 °C, and expression of the target protein was induced at an optical density (A600) of 0.6-0.8 by the addition of isopropyl β-d-1-thiogalactopyranoside (IPTG) to a final concentration of 0.5 mM. The cell culture was incubated at 18 °C for 12-16 hrs. Bacterial cultures were harvested by centrifugation (8,000 ×g, 10 min, 4 °C) and resuspended in lysis buffer containing 25 mM Tris (pH 8.0), 750 mM NaCl, 2 mM DTT, 0.5 mg/mL lysozyme, 0.5 mM phenylmethylsulfonyl fluoride (PMSF) and 0.02 mg/mL DNase I. Bacterial cells were lysed by alternating sonication (30% amplitude, 1 s on/1 s off) and homogenization using a tissue grinder. The lysed cell suspension was clarified by centrifugation (18,000 ×g, 30 min, 4 °C) and the supernatant was incubated with Ni-NTA resin for over 2 hrs at 4 °C on a rotator. The Ni-NTA resin was thoroughly washed with 20 mM imidazole in washing buffer containing 50 mM Tris (pH 8.0), 150 mM NaCl, 2 mM DTT, and SUMO-M^pro^ protein was eluted with elution buffer containing 50 to 300 mM imidazole, 50 mM Tris (pH 8.0), 150 mM NaCl, 2 mM DTT. Fractions containing SUMO-M^pro^ proteins greater than 90% homogeneity were pooled and subjected to dialysis (two times) against a buffer containing 50 mM Tris (pH 8.0), 150 mM NaCl, 2 mM DTT and 10% glycerol. SUMO protease digestion was carried out at 30 °C for 1 hr to remove SUMO tag. Following digestion, SUMO Protease and SUMO tag were removed by Ni-NTA resin. The purified tag-free SARS-CoV-2 M^pro^ mutant proteins were fast frozen in liquid nitrogen and stored at -80 °C.

### Enzymatic Assays

For measurement of K_M_/V_max_ of SARS-CoV-2 M^pro^ mutants, proteolytic reactions were carried out with optimized concentrations of the mutant proteins and series concentrations of FRET substrate ranging from 0 to 200 µM in 100 μL of reaction buffer containing 20 mM HEPES (pH 6.5), 120 mM NaCl, 0.4 mM EDTA, 4 mM DTT, and 20% glycerol at 30 °C in a BioTek Cytation 5 imaging reader (Agilent) with filters for excitation at 360/40 nm and emission at 460/40 nm. Reactions were monitored every 90 s, and the initial velocity of the proteolytic activity was calculated by linear regression for the first 15 min of the kinetic progress curves. The initial velocity was plotted against the FRET substrate concentrations using the classic Michaelis-Menten equation in Prism 8 software.

For IC_50_ measurements, optimized concentrations of the mutant proteins were incubated with series concentrations of GC-376, PF-00835231 or Nirmatrelvir (PF-07321332) in 100 μL of reaction buffer at 30 °C for 15 min, and the reaction was initiated by adding 10 μM FRET substrate. The reaction was monitored for 1 hr, and the initial velocity was calculated for the first 15 min by linear regression. The IC_50_ was determined by plotting the initial velocity against various concentrations of the compounds using log (inhibitor) vs response-variable slope in Prism 8 software.

For K_i_ measurements, optimized concentrations of the mutant proteins were added to 20 μM FRET substrate with various concentrations of GC-376, PF-00835231 or Nirmatrelvir (PF-07321332) in 200 μL of reaction buffer at 30 °C to initiate the proteolytic reaction. The reaction was monitored for 2 hrs and the initial velocity was calculated for the first 90 min by linear regression. The K_i_ was calculated by plotting the initial velocity against various concentrations of the compounds using Morrison plot (tight binding) in Prism 8 software.

### Differential Scanning Fluorimetry (DSF)

The binding of Nirmatrelvir (PF-07321332) to SARS-CoV-2 mutant proteins was monitored by differential scanning fluorimetry (DSF) using a QuantStudio 5 Real-Time PCR System (Thermo Fisher) as previously described (*8*) with minor modifications. Optimized concentrations of the mutant proteins were mixed with different concentrations of Nirmatrelvir (0.2-200 μM) in 50 μL of reaction buffer and incubated at 30 °C for 1 hr. 1× SYPRO orange (Thermo Fisher) were added and the fluorescence signal was recorded under a temperature gradient ranging from 20 to 95 °C (incremental steps of 0.05 °C s^− 1^). The melting temperature (*T*_m_) was calculated as the mid log of the transition phase from the native to the denatured protein using a Boltzmann model in Protein Thermal Shift Software v1.3. Δ*T*_m_ was calculated by subtracting reference melting temperature of proteins in the presence of DMSO from the *T*_m_ in the presence of compounds. Curve fitting was performed using log (inhibitor) vs Δ*T*_m_-variable slope in Prism (v8) software.

### Generation of Nsp5 S144A and H172Y mutant viruses

To generate recombinant Nsp5 S144A and H172Y mutant viruses, T10484G and C10568T nucleotide substitutions were introduced into the SARS-CoV-2 infectious cDNA subclone plasmid using Q5 site-directed mutagenesis kit (NEB, E0554S) respectively, and then verified by sequencing of the plasmid. Virus recovery was conducted as described previously (*21*). Briefly, viral cDNA fragments were ligated in an equal molar ratio to assemble a full-length genomic cDNA with T4 DNA ligase (NEB, M0202L). The ligated full-length cDNA and a SARS-CoV-2-N plasmid were used for *in vitro* transcription using the T7 mMESSAGE mMACHINE T7 transcription kit (ThermoFisher, AM1344). The transcribed viral RNA and N gene sgRNA were subsequently electroporated into Vero E6-TMPRSS2-T2A-ACE2 cells. These cells were maintained in DMEM containing 2% FBS at 37°C. Culture supernatants were collected at the time when the cytopathic effect was evident. All harvested viral stocks were titrated in Vero E6-TMPRSS2-T2A-ACE2 cells and subjected to sequencing of the Nsp5 coding region to validate the genotypes.

### Growth kinetics and Plaque assay

Vero E6-TMPRSS2-T2A-ACE2 were seeded in 12- or 24-well plates a day prior to infection and inoculated with the designated virus at a multiplicity of infection (MOI) of 0.0001 for 1 hour at 37°C. After 1 hour of incubation, the viral inoculum was removed and replaced with fresh DMEM containing 2% FBS. The culture supernatants were collected at the indicated time points and titrated on Vero E6-TMPRSS2-T2A-ACE2 using a plaque assay. For plaque assay, Vero E6-TMPRSS2-T2A-ACE2 cells were seeded in 6- or 12-well plates a day prior to infection. Each viral stock supernatant was serially diluted and inoculated onto the Vero E6 cells. After 1 hour of incubation at 37°C, the inoculum was removed, and cells were subsequently overlaid with a 1.2% 2× DMEM–agarose mixture. After 48 hours, cells were fixed using 4% formaldehyde for 1 hour and stained using 0.1% crystal violet solution after removal of the agarose overlay. Plaques were photographed, counted, and titers were calculated.

### Antiviral assay

Nirmatrelvir dissolved in DMSO was serially diluted in DMEM as a 6-pt dose-response with 5-fold dilutions between test concentrations, starting at 20 μM final concentration. 50 µL diluted compound was then added to a 96-well cell culture plate in triplicate. Vero E6-TMPRSS2-T2A-ACE2 cells were batch inoculated with each virus at an MOI of 0.0001. 50 µL virus-inoculated cells were then added to the 96-well plate loaded with diluted compound at a density of 2,5000 cells/well in DMEM containing 2uM P-glycoprotein inhibitor, CP-100356, and 10% heat-inactivated FBS. Cells were incubated for 2 days at 37°C and assayed for cell viability using a Celltiter-Glo 2.0 assay (Promega, G9242) on a Promega GloMax Discover microplate reader (Promega, GM3000) following the instruction of the manufacturer.

### M^pro^ crystallization and structure determination

SARS-CoV-2 M^pro^ was diluted to 5 mg/mL in protein buffer (50 mM Tris pH 7.0, 150 mM NaCl, 4 mM DTT). To prepare inhibitor complexes, protein was incubated overnight at 4 °C with either 2mM GC-376 or 2mM Nirmatrelvir. Protein with Nirmatrelvir were supplemented with 4% DMSO to enhance solubility of the compound, and precipitation during incubation was removed by centrifugation prior to crystallization. Since GC-376 is water soluble, no precipitation was observed, and centrifugation was not necessary. Crystals were grown by mixing 1.5 μL of the protein solution with 1.5 μL of the precipitant solution in a hanging-drop vapor-diffusion apparatus. Three precipitant conditions were used for crystal growth: **1**. 25% PEG 3350, 0.1 M K/Na tartrate, and 0.005 M MgCl_2_; **2**. 0.2 M NaCl, 10% 1,6-hexanediol, and 20% PEG MME 2K; **3**. 0.1 M MgCl_2_, 20% PEG 3350, 10% 1,6-hexanediol, 0.1 M HEPES pH 7.5, and 0.1 M Li_2_SO_4_. Crystals were transferred to a cryoprotectant solution and flash-frozen in liquid nitrogen. Cryoprotectant solution was varied based on the crystallization condition as follows: **1**. 27.5% PEG 3350, 0.1 M K/Na tartrate, 0.005 M MgCl_2_, and 15% glycerol; **2**. 0.2 M NaCl, 10% 1,6-hexanediol, 20% PEG MME 2K, and 20% glycerol; **3**. 0.1 M MgCl_2_, 15% PEG 3350, 10% 1,6-hexanediol, 0.1 M HEPES pH 7.5, 0.1 M LiSO_4_, and 20% glycerol.

X-ray diffraction data were collected at the Southeast Regional Collaborative Access Team (SER-CAT) 22-ID and 22-BM beamlines at the Advanced Photon Source (APS) in Argonne, IL, and processed with HKL2000 and CCP4. PHASER was used for molecular replacement using a previously solved SARS-CoV-2 M^pro^ structure (PDB ID: 7LYH) as a reference model. The CCP4 suite (*22*), Coot (*23*), and the PDB REDO server (pdb-redo.edu) (*24*) were used to complete the model building and refinement. The PyMOL Molecular Graphics System (Schrödinger, LLC) was used to generate all images.

## SUPPLEMENTARY MATERIALS

Figs. S1-S5

Tables S1-S2

## Author contributions

Conceptualization: J.W., Y.C., Y.H., H.T., E.M.L.; Protein expression: Y.H., H.T., X.Z.; Enzymatic assay and thermal shift assay: Y.H., H.T.; Protein crystallization: E.M.L., R.T.M., M.V.M., Y.C.; Recombinant SARS-CoV-2 assays: X.Z., X.D.; Structure determination: E.M.L., L.M.C.J., S.G.B., Y.C.; Writing: J.W., Y.C., E.M.L.

## Data and materials availability

The X-ray crystal structures have been deposited into the Protein Data Bank with accession codes 8D4J (H172Y apo), 8D4K (H172Y-GC376), 8D4L (S144A), 8D4M (S144A-GC376), 8D4N (E166Q apo), 8DFN (H164N apo), 8DD1 (H164N-GC376), 8DFE (S144L apo), 8DD9 (S144L-GC376), 8DGB (Q192T-GC376), and 8DCZ (M165Y-nirmatrelvir).

