## Supplementary Information for "Naturally occurring mutations of SARS-CoV-2 main protease confer drug resistance to nirmatrelvir"

**This PDF file includes:**

Figs. 1-5

Tables 1-2

#### Supplementary Figures

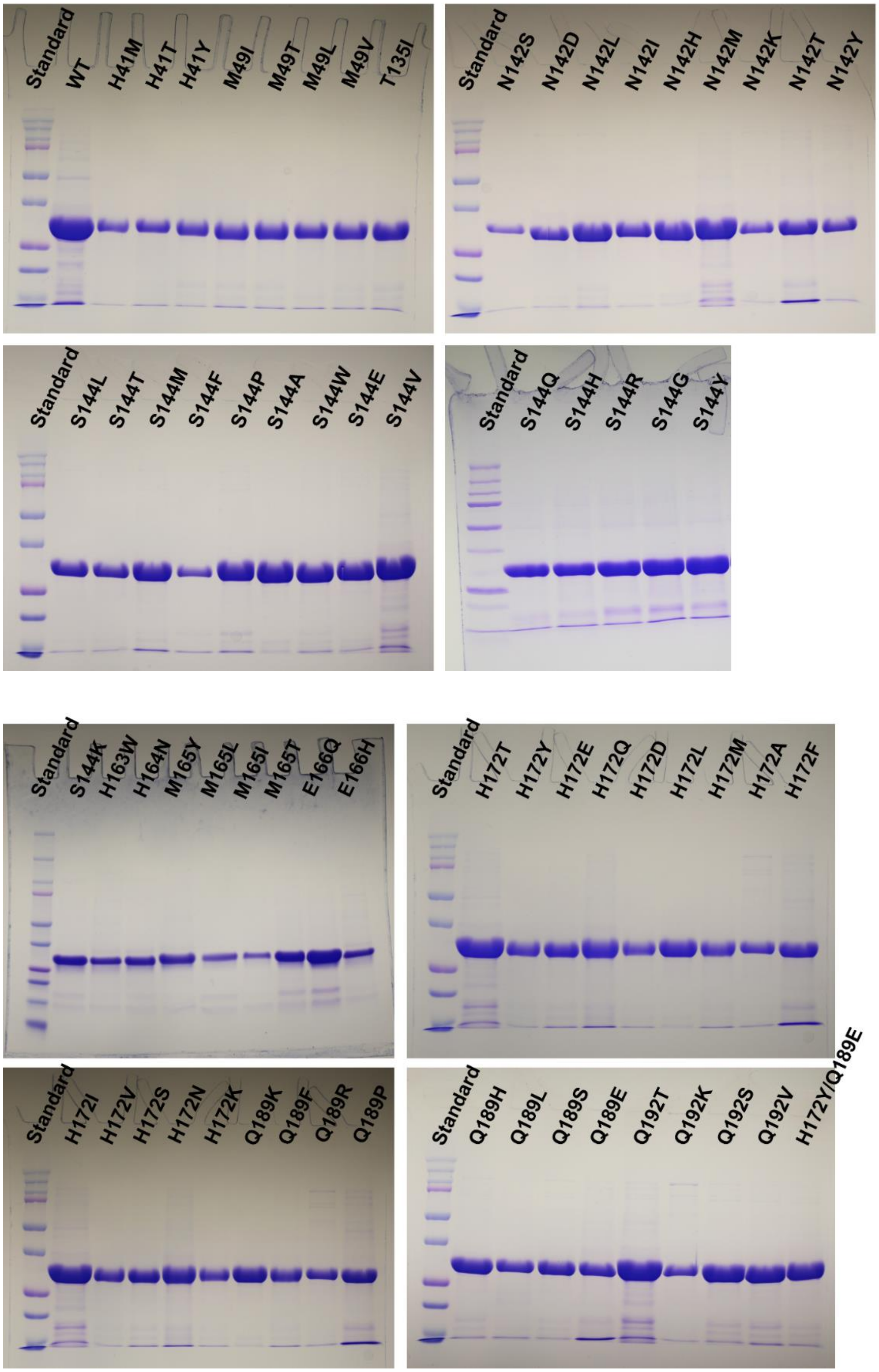

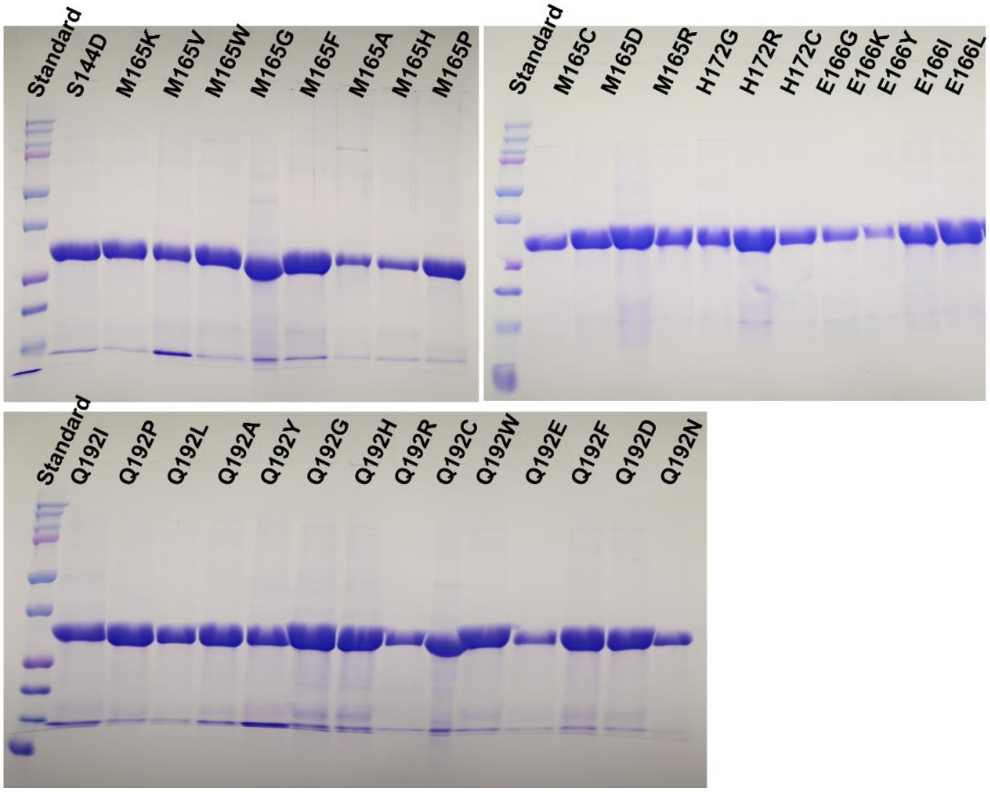

**Supplementary Figure 1. Sodium dodecyl sulfate-polyacrylamide gel electrophoresis (SDS-PAGE) analysis of purified SARS-CoV-2 M$^{pro}$ WT and mutant proteins.** 10 µl of purified proteins were analyzed on 15% SDS-PAGE gel and the protein bands were visualized by staining with Coomassie blue. Protein standard (10-250 kD) was purchased from Bio-Rad, Cat #1610374.

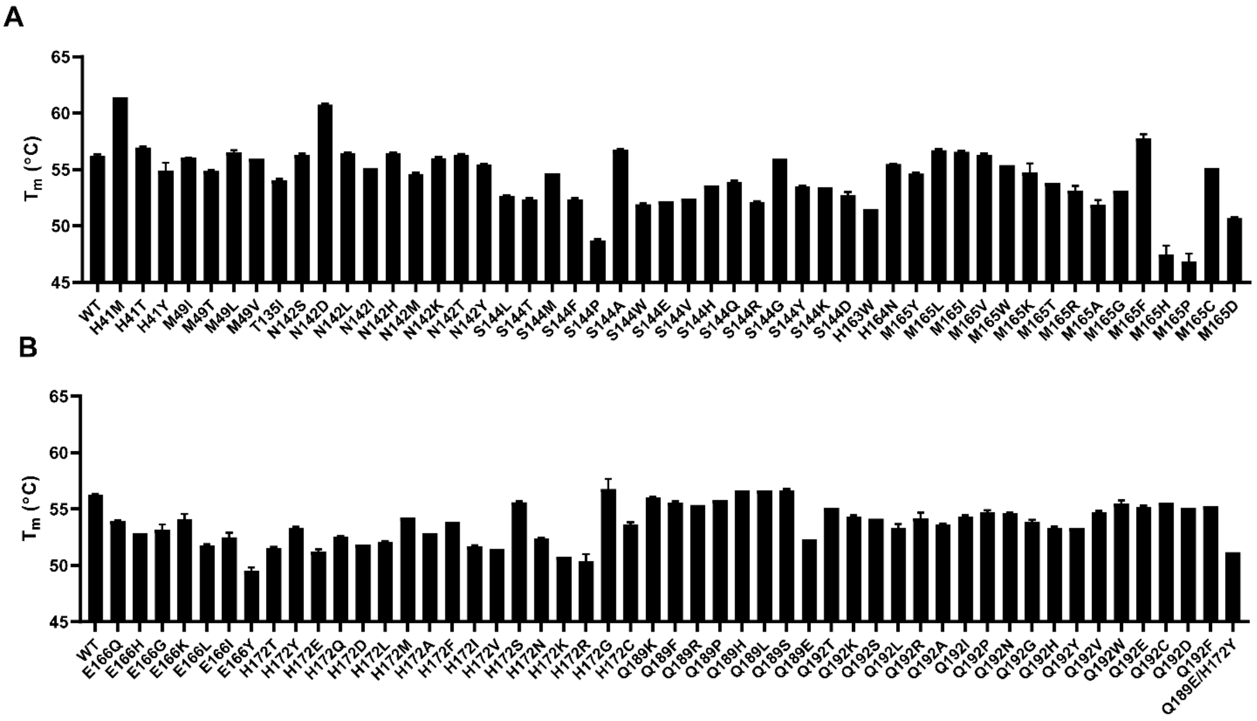

**Supplementary Figure 2.** Thermal shift assay results of $M^{pro}$ mutants.

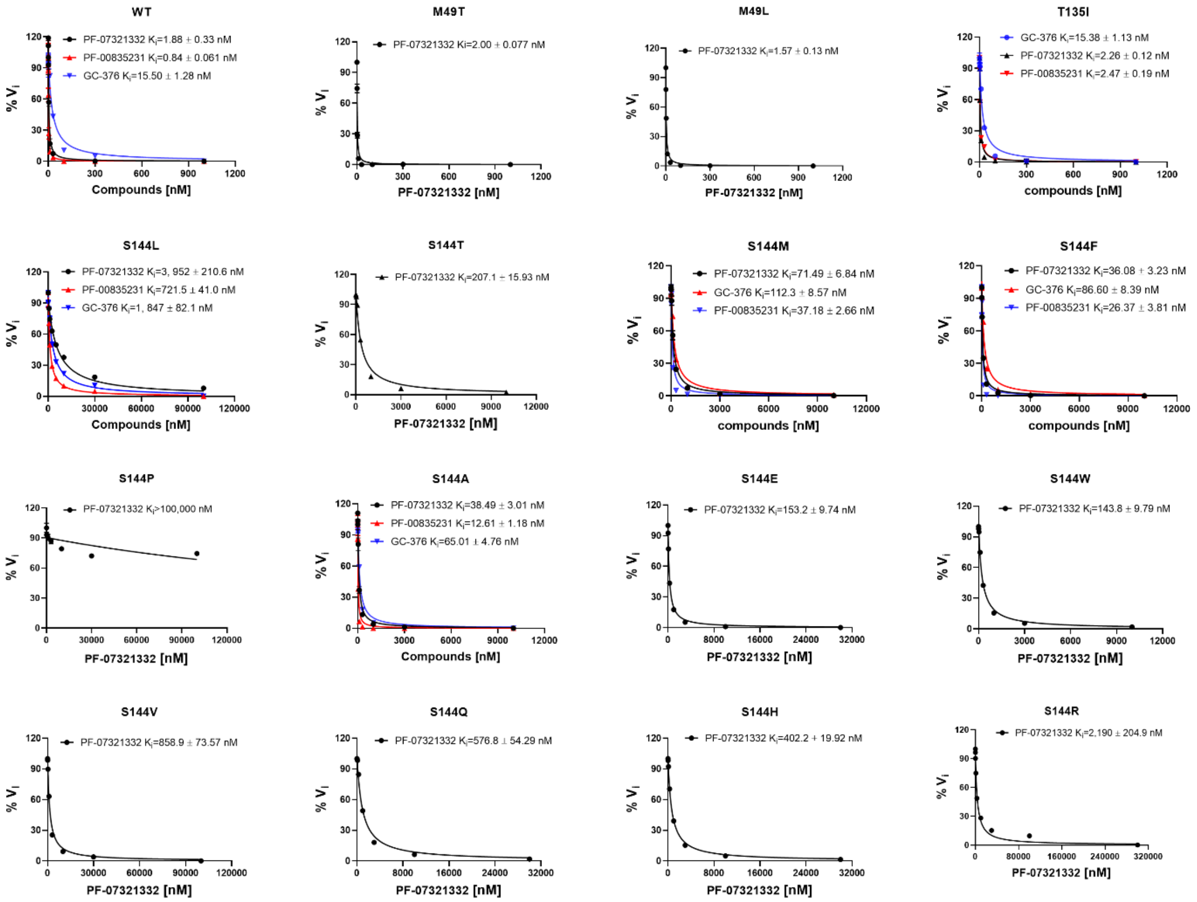
WT
PF-07321332 Ki=1.88 ± 0.33 nM
PF-00835231 Ki=0.84 ± 0.061 nM
GC-376 Ki=15.50 ± 1.28 nM
% Vi
Compounds [nM]
M49T
PF-07321332 Ki=2.00 ± 0.077 nM
% Vi
PF-07321332 [nM]
M49L
PF-07321332 Ki=1.57 ± 0.13 nM
% Vi
PF-07321332 [nM]
T135I
GC-376 Ki=15.38 ± 1.13 nM
PF-07321332 Ki=2.26 ± 0.12 nM
PF-00835231 Ki=2.47 ± 0.19 nM
% Vi
compounds [nM]
S144L
PF-07321332 Ki=3, 952 ± 210.6 nM
PF-00835231 Ki=721.5 ± 41.0 nM
GC-376 Ki=1, 847 ± 82.1 nM
% Vi
Compounds [nM]
S144T
PF-07321332 Ki=207.1 ± 15.93 nM
% Vi
PF-07321332 [nM]
S144M
PF-07321332 Ki=71.49 ± 6.84 nM
GC-376 Ki=112.3 ± 8.57 nM
PF-00835231 Ki=37.18 ± 2.66 nM
% Vi
compounds [nM]
S144F
PF-07321332 Ki=36.08 ± 3.23 nM
GC-376 Ki=86.60 ± 8.39 nM
PF-00835231 Ki=26.37 ± 3.81 nM
% Vi
compounds [nM]
S144P
PF-07321332 Ki>100,000 nM
% Vi
PF-07321332 [nM]
S144A
PF-07321332 Ki=38.49 ± 3.01 nM
PF-00835231 Ki=12.61 ± 1.18 nM
GC-376 Ki=65.01 ± 4.76 nM
% Vi
Compounds [nM]
S144E
PF-07321332 Ki=153.2 ± 9.74 nM
% Vi
PF-07321332 [nM]
S144W
PF-07321332 Ki=143.8 ± 9.79 nM
% Vi
PF-07321332 [nM]
S144V
PF-07321332 Ki=858.9 ± 73.57 nM
% Vi
PF-07321332 [nM]
S144Q
PF-07321332 Ki=576.8 ± 54.29 nM
% Vi
PF-07321332 [nM]
S144H
PF-07321332 Ki=402.2 + 19.92 nM
% Vi
PF-07321332 [nM]
S144R
PF-07321332 Ki=2,190 ± 204.9 nM
% Vi
PF-07321332 [nM]

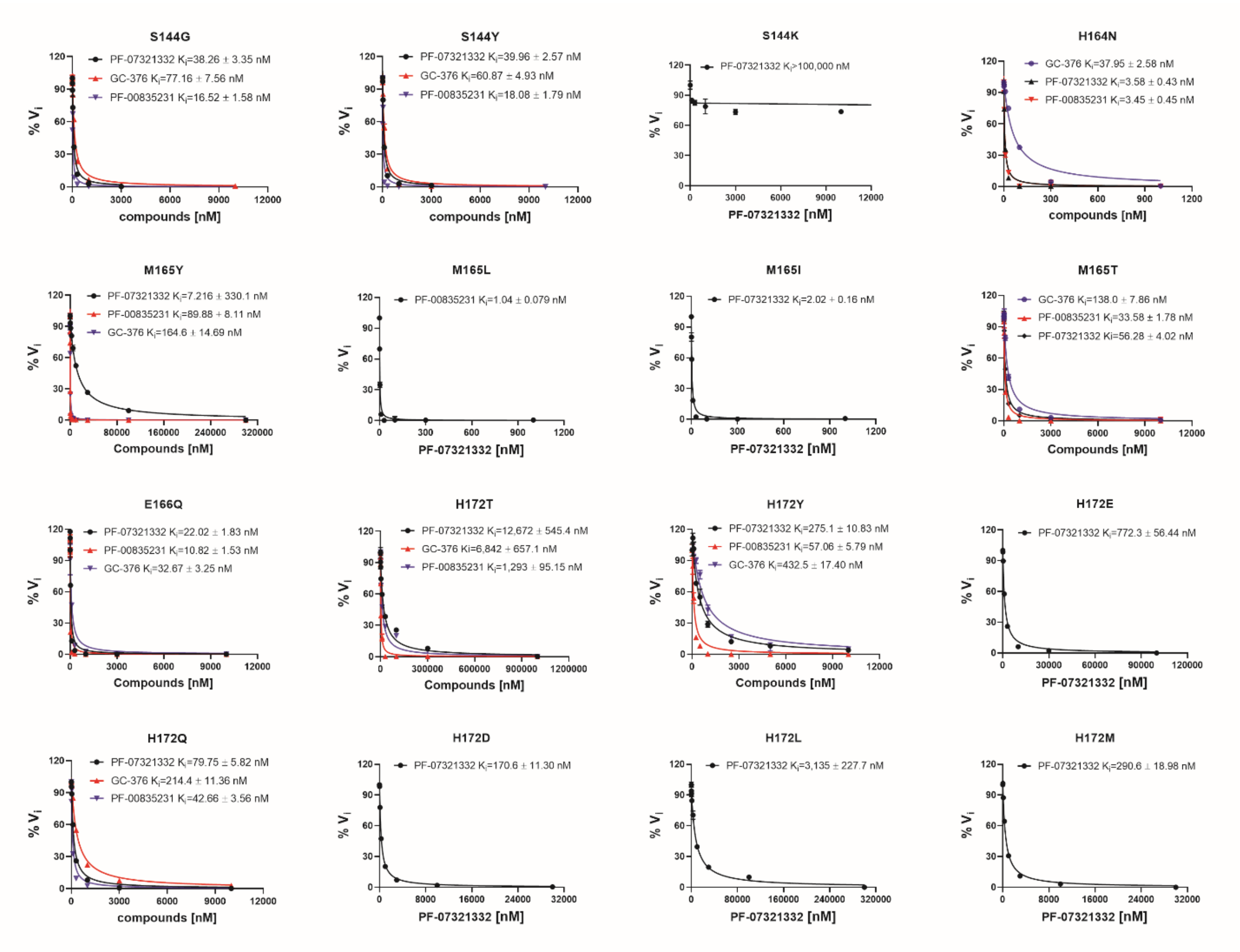
S144G
PF-07321332 Ki=38.26 ± 3.35 nM
GC-376 Ki=77.16 ± 7.56 nM
PF-00835231 Ki=16.52 ± 1.58 nM
% Vi
compounds [nM]
S144Y
PF-07321332 Ki=39.96 ± 2.57 nM
GC-376 Ki=60.87 ± 4.93 nM
PF-00835231 Ki=18.08 ± 1.79 nM
% Vi
compounds [nM]
S144K
PF-07321332 Ki>100,000 nM
% Vi
PF-07321332 [nM]
H164N
GC-376 Ki=37.95 ± 2.58 nM
PF-07321332 Ki=3.58 ± 0.43 nM
PF-00835231 Ki=3.45 ± 0.45 nM
% Vi
compounds [nM]
M165Y
PF-07321332 Ki=7.216 ± 330.1 nM
PF-00835231 Ki=89.88 ± 8.11 nM
GC-376 Ki=164.6 ± 14.69 nM
% Vi
Compounds [nM]
M165L
PF-00835231 Ki=1.04 ± 0.079 nM
% Vi
PF-07321332 [nM]
M165I
PF-07321332 Ki=2.02 ± 0.16 nM
% Vi
PF-07321332 [nM]
M165T
GC-376 Ki=138.0 ± 7.86 nM
PF-00835231 Ki=33.58 ± 1.78 nM
PF-07321332 Ki=56.28 ± 4.02 nM
% Vi
Compounds [nM]
E166Q
PF-07321332 Ki=22.02 ± 1.83 nM
PF-00835231 Ki=10.82 ± 1.53 nM
GC-376 Ki=32.67 ± 3.25 nM
% Vi
Compounds [nM]
H172T
PF-07321332 Ki=12,672 ± 545.4 nM
GC-376 Ki=6,842 ± 657.1 nM
PF-00835231 Ki=1,293 ± 95.15 nM
% Vi
Compounds [nM]
H172Y
PF-07321332 Ki=275.1 ± 10.83 nM
PF-00835231 Ki=57.06 ± 5.79 nM
GC-376 Ki=432.5 ± 17.40 nM
% Vi
Compounds [nM]
H172E
PF-07321332 Ki=772.3 ± 56.44 nM
% Vi
PF-07321332 [nM]
H172Q
PF-07321332 Ki=79.75 ± 5.82 nM
GC-376 Ki=214.4 ± 11.36 nM
PF-00835231 Ki=42.66 ± 3.56 nM
% Vi
compounds [nM]
H172D
PF-07321332 Ki=170.6 ± 11.30 nM
% Vi
PF-07321332 [nM]
H172L
PF-07321332 Ki=3,135 ± 227.7 nM
% Vi
PF-07321332 [nM]
H172M
PF-07321332 Ki=290.6 ± 18.98 nM
% Vi
PF-07321332 [nM]

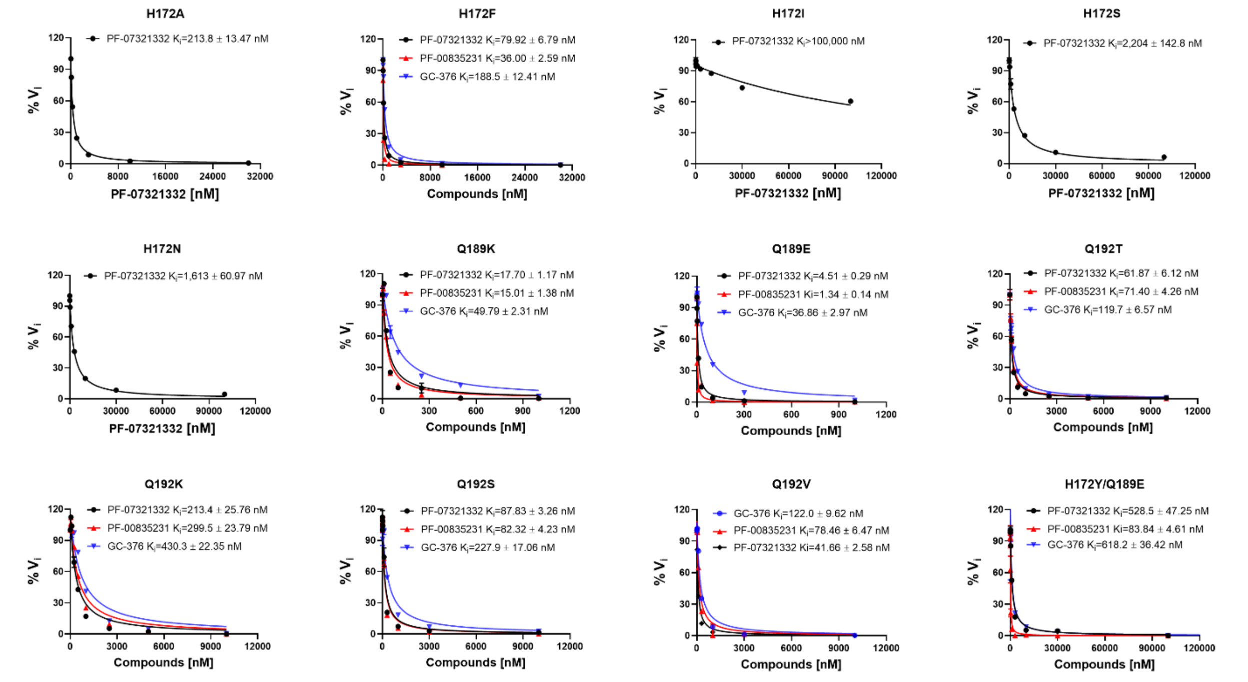
H172A
PF-07321332 Ki=213.8 ± 13.47 nM
% Vi
PF-07321332 [nM]
H172F
PF-07321332 Ki=79.92 ± 6.79 nM
PF-00835231 Ki=36.00 ± 2.59 nM
GC-376 Ki=188.5 ± 12.41 nM
% Vi
Compounds [nM]
H172I
PF-07321332 Ki>100,000 nM
% Vi
PF-07321332 [nM]
H172S
PF-07321332 Ki=2,204 ± 142.8 nM
% Vi
PF-07321332 [nM]
H172N
PF-07321332 Ki=1,613 ± 60.97 nM
% Vi
PF-07321332 [nM]
Q189K
PF-07321332 Ki=17.70 ± 1.17 nM
PF-00835231 Ki=15.01 ± 1.38 nM
GC-376 Ki=49.79 ± 2.31 nM
% Vi
Compounds [nM]
Q189E
PF-07321332 Ki=4.51 ± 0.29 nM
PF-00835231 Ki=1.34 ± 0.14 nM
GC-376 Ki=36.86 ± 2.97 nM
% Vi
Compounds [nM]
Q192T
PF-07321332 Ki=61.87 ± 6.12 nM
PF-00835231 Ki=71.40 ± 4.26 nM
GC-376 Ki=119.7 ± 6.57 nM
% Vi
Compounds [nM]
Q192K
PF-07321332 Ki=213.4 ± 25.76 nM
PF-00835231 Ki=299.5 ± 23.79 nM
GC-376 Ki=430.3 ± 22.35 nM
% Vi
Compounds [nM]
Q192S
PF-07321332 Ki=87.83 ± 3.26 nM
PF-00835231 Ki=82.32 ± 4.23 nM
GC-376 Ki=227.9 ± 17.06 nM
% Vi
Compounds [nM]
Q192V
GC-376 Ki=122.0 ± 9.62 nM
PF-00835231 Ki=78.46 ± 6.47 nM
PF-07321332 Ki=41.66 ± 2.58 nM
% Vi
Compounds [nM]
H172Y/Q189E
PF-07321332 Ki=528.5 ± 47.25 nM
PF-00835231 Ki=83.84 ± 4.61 nM
GC-376 Ki=618.2 ± 36.42 nM
% Vi
Compounds [nM]

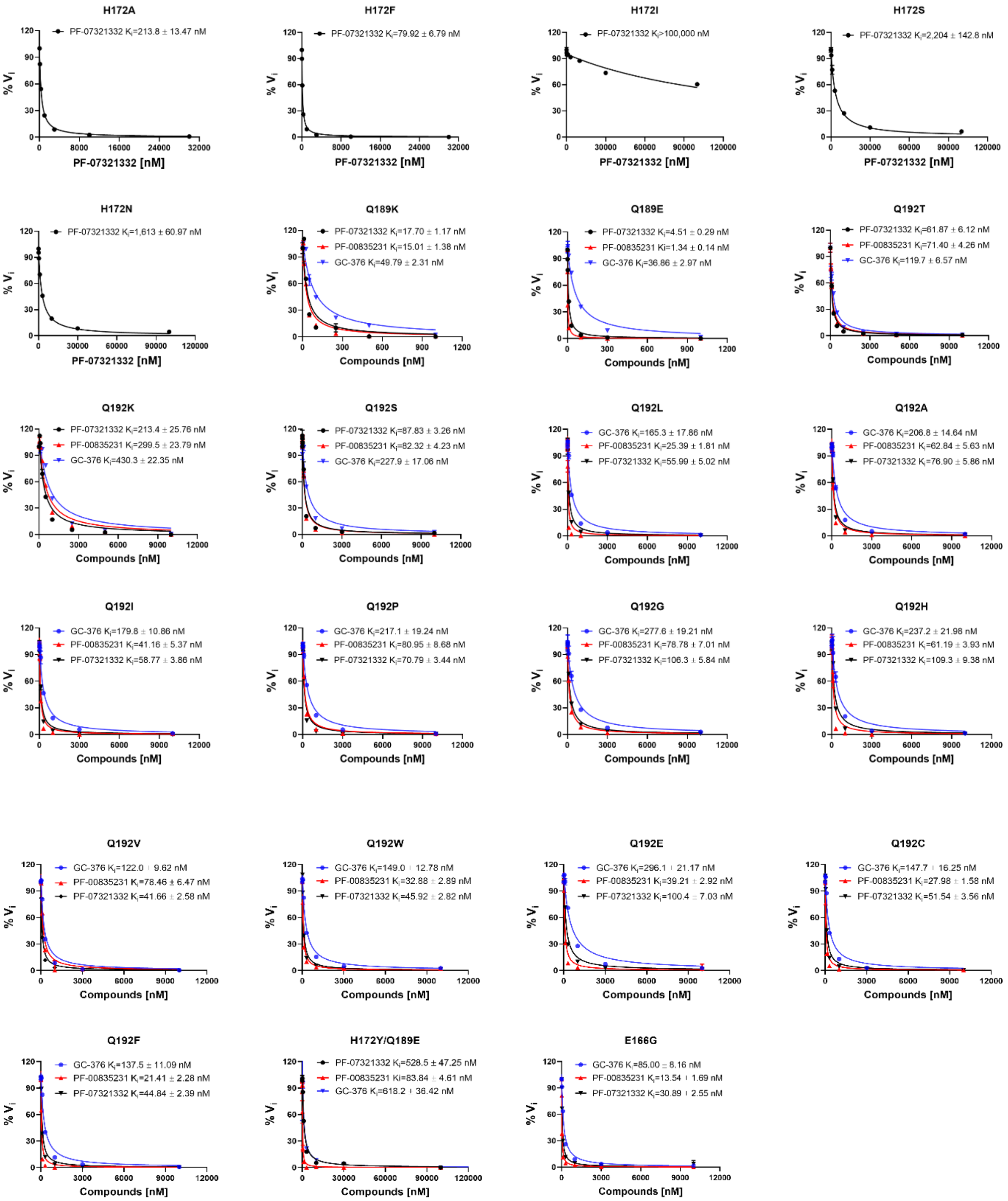

**Supplementary Figure 3.** **Determination of $K_i$ values for GC-376, PF-00835231 or Nirmatrelvir (PF-07321332) against SARS-CoV-2 WT and mutant proteins in FRET assay.** Curves were generated by fitting the initial velocity against various concentrations of the compounds using Morrison plot (tight binding) in Prism 8 software. The results were the average of duplicates.

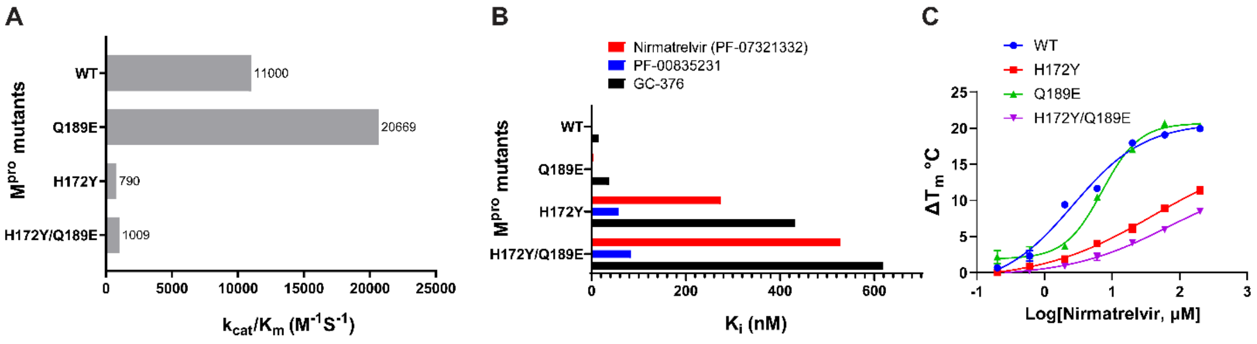

**Supplementary Figure 4. Characterization of the enzymatic activity and drug inhibition of the SARS-CoV-2 $M^{pro}$ H172Y/Q189E double mutant.** (**A**) Plot of the $k_{cat}/K_m$ values for the WT, Q189E, H172Y, and the H172Y/Q189E double mutant. (**B**) Plot of the $K_i$ values for the WT, Q189E, H172Y, and the H172Y/Q189E double mutant. (**C**) Plot of the thermal shift binding assay for the WT, Q189E, H172Y, and the H172Y/Q189E double mutant with nirmatrelvir.

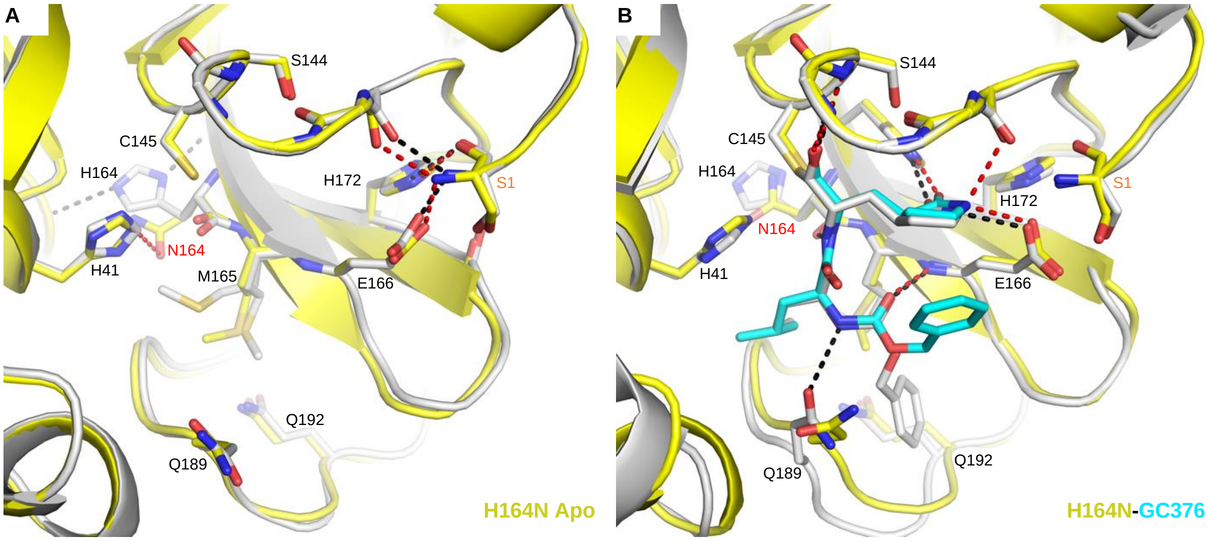

**Supplemental Figure 5. X-ray crystal structures of H164N mutant.** (**A**) Apo M^pro WT (white, PDB 7JP1) aligned with apo M^pro H164N (yellow, PDB 8DFN). (**B**) M^pro WT GC376 complex (white, PDB 6WTT) aligned with M^pro H164N GC376 complex (yellow, PDB 8DD1). WT hydrogen bonds are shown as black dashes, and mutant hydrogen bonds are shown as red dashes. GC-376 is shown in white for the WT structure and cyan for the mutant structure. Mutation is indicated with red text. Ser1 from an adjacent monomer is indicated with orange text.

#### Supplementary Tables

**Table S1. Enzymatic characterization, drug inhibition, and PDB code of SARS-CoV-2 $M^{pro}$ mutants.**

| | | | | | | |
|---|---|---|---|---|---|---|
| **Resistant mutants identified from the GISAID SARS-CoV-2 sequence analysis** | | | | | | |
| $M^{pro}$ mutants | Occurrence[a] | $k_{cat}$, $V_{max}$, $K_m$, $k_{cat}/K_m$ | 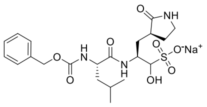 GC-376 | 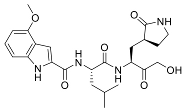 PF-00835231 | 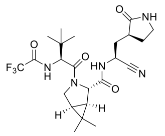 Nirmatrelvir (PF-07321332) | PDB code |
| WT | | $K_m$ = 35.36 ± 2.41 µM $V_{max}$ = 38.97 ± 0.91 nM/s $k_{cat}$ = 0.39 $S^{-1}$ $k_{cat}/K_m$ = 11,000 $S^{-1}M^{-1}$ | $IC_{50}$ = 40.25 ± 1.61 nM $K_i$ = 15.50 ± 1.28 nM | $IC_{50}$ = 15.09 ± 0.80 nM $K_i$ = 0.84 ± 0.061 nM | $IC_{50}$ = 26.03 ± 1.65 nM $K_i$ =1.88 ± 0.33 nM | |
| H41M | 84 | Enzymatically inactive | | | | |
| H41T | 27 | Enzymatically inactive | | | | |
| H41Y | 19 | Enzymatically inactive | | | | |
| M49I | 2,071 | $K_m$ = 21.43 ± 3.29 µM $V_{max}$ = 19.90 ± 0.94 nM/s $k_{cat}$ = 0.3980 $S^{-1}$ $k_{cat}/K_m$ = 18,572 $S^{-1}M^{-1}$ | $IC_{50}$ = 41.55 ± 0.75 nM | $IC_{50}$ = 17.84 ± 0.64 nM | $IC_{50}$ = 26.54 ± 1.05 nM | |
| M49T | 78 | $K_m$ = 21.58 ± 2.61 µM $V_{max}$ = 14.47 ± 1.08 nM/s $k_{cat}$ = 0.2315 $S^{-1}$ $k_{cat}/K_m$ = 10,728 $S^{-1}M^{-1}$ | | | $IC_{50}$ = 15.76 ± 0.69 nM $K_i$ = 2.00 ± 0.08 nM | |
| M49L | 73 | $K_m$ = 29.02 ± 2.42 µM $V_{max}$ = 34.77 ± 1.93 nM/s $k_{cat}$ = 0.5563 $S^{-1}$ $k_{cat}/K_m$ = 19,170 $S^{-1}M^{-1}$ | | | $IC_{50}$ = 18.50 ± 0.69 nM $K_i$ = 1.57 ± 0.13 nM | |
| M49V | 55 | $K_m$ = 32.35 ± 1.65 µM $V_{max}$ = 23.75 ± 0.92 nM/s $k_{cat}$ = 0.1900 $S^{-1}$ $k_{cat}/K_m$ = 5,873 $S^{-1}M^{-1}$ | | | $IC_{50}$ = 23.48 ± 0.39 nM | |
| T135I | 1,340 | $K_m$ = 27.08 ± 2.76 µM $V_{max}$ = 29.28 ± 0.97 nM/s $k_{cat}$ = 0.2928 $S^{-1}$ $k_{cat}/K_m$ = 10,812 $S^{-1}M^{-1}$ | $IC_{50}$ = 42.12 ± 3.41 nM $K_i$ = 15.38 ± 1.13 nM | $IC_{50}$ = 15.63 ± 1.05 nM $K_i$ = 2.47 ± 0.19 nM | $IC_{50}$ = 21.86 ± 0.85 nM $K_i$ = 2.26 ± 0.12 nM | |
| N142S | 141 | $K_m$ = 22.24 ± 3.09 µM $V_{max}$ = 26.07 ± 1.13 nM/s $k_{cat}$ = 0.2607 $S^{-1}$ $k_{cat}/K_m$ = 11,722 $S^{-1}M^{-1}$ | $IC_{50}$ = 39.92 ± 1.36 nM | $IC_{50}$ = 17.64 ± 1.18 nM | $IC_{50}$ = 25.97 ± 1.14 nM | |

| | | | | | | |
|---|---|---|---|---|---|---|
| N142D | 76 | $K_m$ = 29.03 ± 2.98 µM $V_{max}$ = 9.91 ± 0.34 nM/s $k_{cat}$ = 0.0793 $S^{-1}$ $k_{cat}/K_m$ = 2,731 $S^{-1}M^{-1}$ | $IC_{50}$ = 37.90 ± 1.67 nM | $IC_{50}$ = 18.23 ± 0.49 nM | $IC_{50}$ = 24.30 ± 1.46 nM | |
| N142L | 34 | $K_m$ = 25.93 ± 2.51 µM $V_{max}$ = 24.94 ± 1.57 nM/s $k_{cat}$ = 0.3990 $S^{-1}$ $k_{cat}/K_m$ = 15,389 $S^{-1}M^{-1}$ | | | $IC_{50}$ = 15.29 ± 0.42 nM | |
| N142I | 18 | $K_m$ = 31.02 ± 5.85 µM $V_{max}$ = 32.86 ± 2.10 nM/s $k_{cat}$ = 0.5258 $S^{-1}$ $k_{cat}/K_m$ = 16,949 $S^{-1}M^{-1}$ | | | $IC_{50}$ = 17.11 ± 0.82 nM | |
| N142H | 12 | $K_m$ = 41.81 ± 5.14 µM $V_{max}$ = 29.84 ± 1.35 nM/s $k_{cat}$ = 0.2387 $S^{-1}$ $k_{cat}/K_m$ = 5,710 $S^{-1}M^{-1}$ | | | $IC_{50}$ = 19.65 ± 0.32 nM | |
| N142M | 9 | $K_m$ = 10.02 ± 3.40 µM $V_{max}$ = 10.88 ± 0.96 nM/s $k_{cat}$ = 0.1741 $S^{-1}$ $k_{cat}/K_m$ = 17,373 $S^{-1}M^{-1}$ | | | $IC_{50}$ = 19.01 ± 0.62 nM | |
| N142K | 8 | $K_m$ = 43.57 ± 4.72 µM $V_{max}$ = 20.14 ± 0.81 nM/s $k_{cat}$ = 0.3222 $S^{-1}$ $k_{cat}/K_m$ = 7,396 $S^{-1}M^{-1}$ | | | $IC_{50}$ = 13.93 ± 0.39 nM | |
| N142T | 4 | $K_m$ = 21.99 ± 4.78 µM $V_{max}$ = 13.66 ± 0.92 nM/s $k_{cat}$ = 0.2186 $S^{-1}$ $k_{cat}/K_m$ = 9,939 $S^{-1}M^{-1}$ | | | $IC_{50}$ = 18.73 ± 0.96 nM | |
| N142Y | 4 | $K_m$ = 20.84 ± 2.45 µM $V_{max}$ = 11.95 ± 0.86 nM/s $k_{cat}$ = 0.1912 $S^{-1}$ $k_{cat}/K_m$ = 9,175 $S^{-1}M^{-1}$ | | | $IC_{50}$ = 14.90 ± 1.26 nM | |
| S144L | 51 | $K_m$ = 51.29 ± 3.20 µM $V_{max}$ = 6.11 ± 0.15 nM/s $k_{cat}$ = 0.0031 $S^{-1}$ $k_{cat}/K_m$ = 60 $S^{-1}M^{-1}$ | $IC_{50}$ = 1,812 ± 86.1 nM $K_i$ = 1,847 ± 82.1 nM | $IC_{50}$ = 716.7 ± 42.5 nM $K_i$ = 721.5 ± 41.0 nM | $IC_{50}$ = 5,364 ± 498.6 nM $K_i$ = 3,952 ± 210.6 nM | Apo: 8DFE GC-376: 8DD9 |
| S144T | 17 | $K_m$ = 44.31 ± 2.92 µM $V_{max}$ = 16.85 ± 0.42 nM/s $k_{cat}$ = 0.0225 $S^{-1}$ $k_{cat}/K_m$ = 507 $S^{-1}M^{-1}$ | | | $IC_{50}$ = 353.7 ± 14.36 nM $K_i$ = 207.1 ± 15.93 nM | |
| S144M | 16 | $K_m$ = 51.85 ± 3.37 µM $V_{max}$ = 28.36 ± 0.72 nM/s $k_{cat}$ = 0.0709 $S^{-1}$ $k_{cat}/K_m$ = 1367 $S^{-1}M^{-1}$ | $K_i$ = 112.3 ± 8.57 nM | $K_i$ = 37.18 ± 2.66 nM | $IC_{50}$ = 175.2 ± 9.72 nM $K_i$ = 79.26 ± 4.13 nM | |
| S144F | 15 | $K_m$ = 45.11 ± 3.45 µM $V_{max}$ = 21.22 ± 0.61 nM/s $k_{cat}$ = 0.0849 $S^{-1}$ | $K_i$ = 86.60 ± 8.39 nM | $K_i$ = 26.37 ± 3.81 nM | $IC_{50}$ = 133.3 ± 5.61 nM $K_i$ = 47.23 ± 2.24 nM | |

| | | | | | | |
|---|---|---|---|---|---|---|
| | | $k_{cat}/K_m$ = 1,882 $S^{-1}M^{-1}$ | | | | |
| S144P | 12 | $K_m$ = 2.78 ± 0.58 µM<br>$V_{max}$ = 0.58 ± 0.024 nM/s<br>$k_{cat}$ = 0.000058 $S^{-1}$<br>$k_{cat}/K_m$ = 21 $S^{-1}M^{-1}$ | | | $IC_{50}$ > 10 µM<br>$K_i$ > 10 µM | |
| S144A | 9 | $K_m$ = 25.66 ± 1.76 µM<br>$V_{max}$ = 38.95 ± 1.12 nM/s<br>$k_{cat}$ = 0.1558 $S^{-1}$<br>$k_{cat}/K_m$ = 6,072 $S^{-1}M^{-1}$ | $IC_{50}$ = 139.80 ± 9.29 nM<br>$K_i$ = 65.01 ± 4.76 nM | $IC_{50}$ = 39.43 ± 1.14 nM<br>$K_i$ = 12.61 ± 1.18 nM | $IC_{50}$ = 171.1 ± 5.33 nM<br>$K_i$ = 36.43 ± 4.11 nM | Apo: 8D4L<br>GC-376: 8D4M |
| S144W | 7 | $K_m$ = 56.59 ± 3.01 µM<br>$V_{max}$ = 15.55 ± 0.39 nM/s<br>$k_{cat}$ = 0.0311 $S^{-1}$<br>$k_{cat}/K_m$ = 550 $S^{-1}M^{-1}$ | | | $IC_{50}$ = 289.60 ± 14.78 nM<br>$K_i$ = 143.80 ± 9.79 nM | |
| S144E | 6 | $K_m$ = 52.69 ± 3.94 µM<br>$V_{max}$ = 23.31 ± 0.69 nM/s<br>$k_{cat}$ = 0.0233 $S^{-1}$<br>$k_{cat}/K_m$ = 442 $S^{-1}M^{-1}$ | | | $IC_{50}$ = 423.40 ± 29.31 nM<br>$K_i$ = 153.20 ± 9.74 nM | |
| S144V | 4 | $K_m$ = 54.43 ± 3.58 µM<br>$V_{max}$ = 17.12 ± 0.45 nM/s<br>$k_{cat}$ = 0.0086 $S^{-1}$<br>$k_{cat}/K_m$ = 157 $S^{-1}M^{-1}$ | | | $IC_{50}$ = 1,442 ± 127.2 nM<br>$K_i$ = 858.9 ± 73.57 nM | |
| S144H | 4 | $K_m$ = 60.72 ± 4.26 µM<br>$V_{max}$ = 16.45 ± 0.48 nM/s<br>$k_{cat}$ = 0.0329 $S^{-1}$<br>$k_{cat}/K_m$ = 542 $S^{-1}M^{-1}$ | | | $IC_{50}$ = 650.5 ± 45.5 nM<br>$K_i$ = 402.2 ± 19.92 nM | |
| S144Q | 3 | $K_m$ = 64.17 ± 4.49 µM<br>$V_{max}$ = 22.19 ± 0.65 nM/s<br>$k_{cat}$ = 0.0222 $S^{-1}$<br>$k_{cat}/K_m$ = 346 $S^{-1}M^{-1}$ | | | $IC_{50}$ = 831.2 ± 48.6 nM<br>$K_i$ = 576.8 ± 54.29 nM | |
| S144R | 2 | $K_m$ = 62.54 ± 5.09 µM<br>$V_{max}$ = 11.57 ± 0.39 nM/s<br>$k_{cat}$ = 0.0014 $S^{-1}$<br>$k_{cat}/K_m$ = 23 $S^{-1}M^{-1}$ | | | $IC_{50}$ = 6,134 ± 704.8 nM<br>$K_i$ = 2,190 ± 204.9 nM | |
| S144G | 2 | $K_m$ = 53.52 ± 4.72 µM<br>$V_{max}$ = 27.78 ± 0.97 nM/s<br>$k_{cat}$ = 0.2222 $S^{-1}$<br>$k_{cat}/K_m$ = 4,152 $S^{-1}M^{-1}$ | $K_i$ = 77.16 ± 7.56 nM | $K_i$ = 16.52 ± 1.58 nM | $IC_{50}$ = 96.55 ± 1.93 nM<br>$K_i$ = 27.98 ± 2.76 nM | |
| S144Y | 2 | $K_m$ = 55.15 ± 4.39 µM<br>$V_{max}$ = 19.44 ± 0.62 nM/s<br>$k_{cat}$ = 0.0778 $S^{-1}$<br>$k_{cat}/K_m$ = 1,410 $S^{-1}M^{-1}$ | $K_i$ = 60.87 ± 4.93 nM | $K_i$ = 18.08 ± 1.79 nM | $IC_{50}$ = 61.49 ± 3.18 nM<br>$K_i$ = 34.09 ± 3.67 nM | |
| S144K | 2 | $K_m$ = 3.88 ± 0.31 µM<br>$V_{max}$ = 0.80 ± 0.041 nM/s<br>$k_{cat}$ = 0.00008 $S^{-1}$<br>$k_{cat}/K_m$ = 21 $S^{-1}M^{-1}$ | | | $IC_{50}$ > 10 µM<br>$K_i$ > 10 µM | |
| S144D | 1 | $K_m$ = 27.60 ± 2.62 µM<br>$V_{max}$ = 5.91 ± 0.18 nM/s<br>$k_{cat}$ = 0.0035 $S^{-1}$ | | | $IC_{50}$ = 8,532 ± 99.83 nM | |

| | | | | | | |
|---|---|---|---|---|---|---|
| | | $k_{cat}/K_m$ = 128 $S^{-1}M^{-1}$ | | | | |
| H163W | 4,672 | Enzymatically inactive | | | | |
| H164N | 4,681 | $K_m$ = 30.72 ± 2.17 µM<br>$V_{max}$ = 19.88 ± 0.69 nM/s<br>$k_{cat}$ = 0.0795 $S^{-1}$<br>$k_{cat}/K_m$ = 2,588 $S^{-1}M^{-1}$ | $IC_{50}$ = 126.3 ± 6.53 nM<br>$K_i$ = 37.95 ± 2.58 nM | $IC_{50}$ = 16.39 ± 0.92 nM<br>$K_i$ = 3.45 ± 0.45 nM | $IC_{50}$ = 32.85 ± 1.92 nM<br>$K_i$ = 3.58 ± 0.43 nM | Apo: 8DFN<br>GC-376: 8DD1 |
| M165Y | 4,677 | $K_m$ = 38.42 ± 2.32 µM<br>$V_{max}$ = 20.26 ± 0.88 nM/s<br>$k_{cat}$ = 0.0101 $S^{-1}$<br>$k_{cat}/K_m$ = 264 $S^{-1}M^{-1}$ | $IC_{50}$ = 423.0 ± 22.3 nM<br>$K_i$ = 164.6 ± 14.69 nM | $IC_{50}$ = 286.2 ± 15.99 nM<br>$K_i$ = 89.88 ± 8.11 nM | $IC_{50}$ = 10,462 ± 497.7 nM<br>$K_i$ = 7,216 ± 330.1 nM | Nirmatrelvir: 8DCZ |
| M165L | 276 | $K_m$ = 20.60 ± 4.24 µM<br>$V_{max}$ = 17.54 ± 1.10 nM/s<br>$k_{cat}$ = 0.2806 $S^{-1}$<br>$k_{cat}/K_m$ = 13,623 $S^{-1}M^{-1}$ | | | $IC_{50}$ = 22.62 ± 0.77 nM<br>$K_i$ =1.04 ± 0.079 nM | |
| M165I | 101 | $K_m$ = 29.45 ± 4.87 µM<br>$V_{max}$ = 14.90 ± 0.82 nM/s<br>$k_{cat}$ = 0.2384 $S^{-1}$<br>$k_{cat}/K_m$ = 8,095 $S^{-1}M^{-1}$ | | | $IC_{50}$ = 27.38 ± 2.09 nM<br>$K_i$ =2.02 ± 0.16 nM | |
| M165V | 16 | $K_m$ = 36.18 ± 2.61 µM<br>$V_{max}$ = 33.75 ± 1.44 nM/s<br>$k_{cat}$ = 0.3375 $S^{-1}$<br>$k_{cat}/K_m$ = 9,328 $S^{-1}M^{-1}$ | | | $IC_{50}$ = 23.96 ± 0.72 nM | |
| M165W | 11 | $K_m$ = 9.41 ± 0.92 µM<br>$V_{max}$ = 1.34 ± 0.066 nM/s<br>$k_{cat}$ = 0.0001 $S^{-1}$<br>$k_{cat}/K_m$ = 12 $S^{-1}M^{-1}$ | | | $IC_{50}$ > 10 µM | |
| M165K | 8 | $K_m$ = 36.41 ± 2.31 µM<br>$V_{max}$ = 10.09 ± 0.33 nM/s<br>$k_{cat}$ = 0.0012 $S^{-1}$<br>$k_{cat}/K_m$ = 33 $S^{-1}M^{-1}$ | | | $IC_{50}$ >10 µM | |
| M165T | 7 | $K_m$ = 55.37 ± 3.49 µM<br>$V_{max}$ = 29.33 ± 0.74 nM/s<br>$k_{cat}$ = 0.0733 $S^{-1}$<br>$k_{cat}/K_m$ = 1,324 $S^{-1}M^{-1}$ | $IC_{50}$ = 239.6 ± 9.64 nM<br>$K_i$ = 138.0 ± 7.86 nM | $IC_{50}$ = 109.2 ± 4.92 nM<br>$K_i$ = 33.58 ± 1.78 nM | $IC_{50}$ = 94.68 ± 5.94 nM<br>$K_i$ = 52.68 ± 4.68 nM | |
| M165R | 5 | $K_m$ = 7.83 ± 0.76 µM<br>$V_{max}$ = 2.32 ± 0.11 nM/s<br>$k_{cat}$ = 0.0003 $S^{-1}$<br>$k_{cat}/K_m$ = 33 $S^{-1}M^{-1}$ | | | $IC_{50}$ > 10 µM | |
| M165A | 4 | $K_m$ = 50.82 ± 3.76 µM<br>$V_{max}$ = 36.68 ± 1.99 nM/s<br>$k_{cat}$ = 0.3668 $S^{-1}$<br>$k_{cat}/K_m$ = 7,218 $S^{-1}M^{-1}$ | | | $IC_{50}$ = 23.28 ± 1.88 nM | |

| | | | | | | |
|---|---|---|---|---|---|---|
| M165G | 3 | $K_m$ = 56.00 ± 4.91 µM<br>$V_{max}$ = 24.53 ± 0.86 nM/s<br>$k_{cat}$ = 0.0245 $S^{-1}$<br>$k_{cat}/K_m$ = 438 $S^{-1}M^{-1}$ | | | $IC_{50}$ = 243.5 ± 16.40 nM | |
| M165F | 3 | $K_m$ = 74.74 ± 3.47 µM<br>$V_{max}$ = 11.72 ± 0.24 nM/s<br>$k_{cat}$ = 0.0059 $S^{-1}$<br>$k_{cat}/K_m$ = 78 $S^{-1}M^{-1}$ | | | $IC_{50}$ = 1,336 ± 49.13 nM | |
| M165H | 2 | $K_m$ = 14.71 ± 1.92 µM<br>$V_{max}$ = 3.49 ± 0.13 nM/s<br>$k_{cat}$ = 0.0003 $S^{-1}$<br>$k_{cat}/K_m$ = 20 $S^{-1}M^{-1}$ | | | $IC_{50}$ > 10 µM | |
| M165P | 2 | $K_m$ = 0.11 ± 0.024 µM<br>$V_{max}$ = 1.02 ± 0.10 nM/s<br>$k_{cat}$ = 0.0001 $S^{-1}$<br>$k_{cat}/K_m$ = 773 $S^{-1}M^{-1}$ | | | $IC_{50}$ > 10 µM | |
| M165C | 1 | $K_m$ = 46.24 ± 2.54 µM<br>$V_{max}$ = 33.39 ± 0.96 nM/s<br>$k_{cat}$ = 0.3339 $S^{-1}$<br>$k_{cat}/K_m$ = 7,221 $S^{-1}M^{-1}$ | | | $IC_{50}$ = 24.84 ± 1.77 nM | |
| M165D | 1 | $K_m$ = 68.73 ± 4.65 µM<br>$V_{max}$ = 16.03 ± 0.46 nM/s<br>$k_{cat}$ = 0.0178 $S^{-1}$<br>$k_{cat}/K_m$ = 258 $S^{-1}M^{-1}$ | | | $IC_{50}$ = 147.6 ± 10.59 nM | |
| E166Q | 4,681 | $K_m$ = 36.04 ± 3.16 µM<br>$V_{max}$ = 50.18 ± 1.55 nM/s<br>$k_{cat}$ = 0.4014 $S^{-1}$<br>$k_{cat}/K_m$ = 11,139 $S^{-1}M^{-1}$ | $IC_{50}$ = 52.31 ± 2.06 nM<br>$K_i$ = 47.10 ± 3.49 nM | $IC_{50}$ = 14.88 ± 0.47 nM<br>$K_i$ = 4.45 ± 0.39 nM | $IC_{50}$ = 27.87 ± 2.11 nM<br>$K_i$ = 8.37 ± 0.74 nM | Apo: 8D4N |
| E166H | 232 | $K_m$ = 55.96 ± 5.14 µM<br>$V_{max}$ = 17.55 ± 0.65 nM/s<br>$k_{cat}$ = 0.0351 $S^{-1}$<br>$k_{cat}/K_m$ = 627 $S^{-1}M^{-1}$ | | | $IC_{50}$ = 9,598 ± 1,675 nM | |
| E166G | 16 | $K_m$ = 32.90 ± 3.16 µM<br>$V_{max}$ = 24.39 ± 0.98 nM/s<br>$k_{cat}$ = 0.04878 $S^{-1}$<br>$k_{cat}/K_m$ = 1,483 $S^{-1}M^{-1}$ | $K_i$ = 85.00 ± 8.16 nM | $K_i$ = 13.54 ± 1.69 nM | $IC_{50}$ = 92.15 ± 6.49 nM<br>$K_i$ = 30.89 ± 2.55 nM | |
| E166K | 10 | $K_m$ = 50.12 ± 4.68 µM<br>$V_{max}$ = 25.93 ± 1.17 nM/s<br>$k_{cat}$ = 0.02593 $S^{-1}$<br>$k_{cat}/K_m$ = 517 $S^{-1}M^{-1}$ | | | $IC_{50}$ = 1,994 ± 175.1 nM | |
| E166L | 3 | $K_m$ = 18.73 ± 0.99 µM<br>$V_{max}$ = 2.16 ± 0.034 nM/s<br>$k_{cat}$ = 0.000432 $S^{-1}$<br>$k_{cat}/K_m$ = 23 $S^{-1}M^{-1}$ | | | $IC_{50}$ = 214.2 ± 18.57 nM | |
| E166Y | 2 | $K_m$ = 27.92 ± 1.67 µM<br>$V_{max}$ = 11.00 ± 0.43 nM/s<br>$k_{cat}$ = 0.011 $S^{-1}$<br>$k_{cat}/K_m$ = 394 $S^{-1}M^{-1}$ | | | $IC_{50}$ = 3,711 ± 283.1 nM | |

| | | | | | | |
|---|---|---|---|---|---|---|
| E166I | 1 | $K_m$ = 37.82 ± 3.77 µM<br>$V_{max}$ = 2.00 ± 0.14 nM/s<br>$k_{cat}$ = 0.0002 $S^{-1}$<br>$k_{cat}/K_m$ = 5 $S^{-1}M^{-1}$ | | | $IC_{50}$ > 10,000 nM | |
| H172T | 96 | $K_m$ = 51.84 ± 3.01 µM<br>$V_{max}$ = 6.29 ± 0.14 nM/s<br>$k_{cat}$ = 0.0006 $S^{-1}$<br>$k_{cat}/K_m$ = 12 $S^{-1}M^{-1}$ | $IC_{50}$ > 10,000 nM<br>$K_i$ = 6,842 ± 657.1 nM | $IC_{50}$ = 7,888 ± 420 nM<br>$K_i$ = 1,293 ± 95.15 nM | $IC_{50}$ > 10,000 nM<br>$K_i$ = 12,672 ± 545.4 nM | |
| H172Y | 21 | $K_m$ = 39.20 ± 2.83 µM<br>$V_{max}$ = 15.26 ± 0.39 nM/s<br>$k_{cat}$ = 0.0305 $S^{-1}$<br>$k_{cat}/K_m$ = 790 $S^{-1}M^{-1}$ | $IC_{50}$ = 449.2 ± 27.81 nM<br>$K_i$ = 432.5 ± 17.40 nM | $IC_{50}$ = 126.4 ± 5.64 nM<br>$K_i$ = 57.06 ± 5.79 nM | $IC_{50}$ = 279.3 ± 23.09 nM<br>$K_i$ = 275.1 ± 10.83 nM | Apo: 8D4J<br>GC-376: 8D4K |
| H172E | 16 | $K_m$ = 13.43 ± 0.88 µM<br>$V_{max}$ = 3.23 ± 0.12 nM/s<br>$k_{cat}$ = 0.000323 $S^{-1}$<br>$k_{cat}/K_m$ = 24 $S^{-1}M^{-1}$ | | | $IC_{50}$ > 10,000 nM | |
| H172Q | 12 | $K_m$ = 49.61 ± 4.41 µM<br>$V_{max}$ = 21.53 ± 0.74 nM/s<br>$k_{cat}$ = 0.1722 $S^{-1}$<br>$k_{cat}/K_m$ = 3,472 $S^{-1}M^{-1}$ | $K_i$ = 214.4 ± 11.36 nM | $K_i$ = 42.66 ± 3.56 nM | $IC_{50}$ = 152.1 ± 8.01 nM<br>$K_i$ = 70.54 ± 3.90 nM | |
| H172D | 10 | $K_m$ = 53.01 ± 4.79 µM<br>$V_{max}$ = 13.87 ± 0.50 nM/s<br>$k_{cat}$ = 0.0277 $S^{-1}$<br>$k_{cat}/K_m$ = 523 $S^{-1}M^{-1}$ | | | $IC_{50}$ = 266.1 ± 24.8 nM<br>$K_i$ = 170.6 ± 11.30 nM | |
| H172L | 7 | $K_m$ = 54.89 ± 2.08 µM<br>$V_{max}$ = 5.99 ± 0.11 nM/s<br>$k_{cat}$ = 0.0012 $S^{-1}$<br>$k_{cat}/K_m$ = 22 $S^{-1}M^{-1}$ | | | $IC_{50}$ = 3,380 ± 295.48 nM<br>$K_i$ = 3,135 ± 227.7 nM | |
| H172M | 7 | $K_m$ = 60.05 ± 3.65 µM<br>$V_{max}$ = 19.62 ± 0.49 nM/s<br>$k_{cat}$ = 0.0196 $S^{-1}$<br>$k_{cat}/K_m$ = 327 $S^{-1}M^{-1}$ | | | $IC_{50}$ = 699.0 ± 27.0 nM<br>$K_i$ = 290.6 ± 18.98 nM | |
| H172A | 7 | $K_m$ = 56.15 ± 3.40 µM<br>$V_{max}$ = 27.33 ± 0.67 nM/s<br>$k_{cat}$ = 0.0547 $S^{-1}$<br>$k_{cat}/K_m$ = 973 $S^{-1}M^{-1}$ | | | $IC_{50}$ = 374.7 ± 30.5 nM<br>$K_i$ = 213.8 ± 13.47 nM | |
| H172F | 5 | $K_m$ = 66.53 ± 2.68 µM<br>$V_{max}$ = 29.64 ± 0.51 nM/s<br>$k_{cat}$ = 0.0741 $S^{-1}$<br>$k_{cat}/K_m$ = 1,114 $S^{-1}M^{-1}$ | $K_i$ = 188.5 ± 12.41 nM | $K_i$ = 36.00 ± 2.59 nM | $IC_{50}$ = 212.2 ± 9.69 nM<br>$K_i$ = 46.60 ± 5.59 nM | |
| H172I | 5 | $K_m$ = 61.31 ± 4.27 µM<br>$V_{max}$ = 58.34 ± 1.68 nM/s<br>$k_{cat}$ = 0.00583 $S^{-1}$<br>$k_{cat}/K_m$ = 95 $S^{-1}M^{-1}$ | | | $IC_{50}$ > 10 µM<br>$K_i$ > 10 µM | |
| H172V | 5 | $K_m$ = 37.81 ± 1.79 µM<br>$V_{max}$ = 7.14 ± 0.12 nM/s<br>$k_{cat}$ = 0.0014 $S^{-1}$<br>$k_{cat}/K_m$ = 38 $S^{-1}M^{-1}$ | | | $IC_{50}$ > 10 µM | |

| | | | | | |
|---|---|---|---|---|---|
| H172S | 4 | $K_m$ = 65.48 ± 2.33 µM<br>$V_{max}$ = 15.5 ± 0.23 nM/s<br>$k_{cat}$ = 0.0052 $S^{-1}$<br>$k_{cat}/K_m$ = 79 $S^{-1}M^{-1}$ | | | $IC_{50}$ = 5,650 ± 180.8 nM<br>$K_i$ = 2,204 ± 142.8 nM |
| H172N | 4 | $K_m$ = 62.46 ± 2.12 µM<br>$V_{max}$ = 15.33 ± 0.22 nM/s<br>$k_{cat}$ = 0.0051 $S^{-1}$<br>$k_{cat}/K_m$ = 82 $S^{-1}M^{-1}$ | | | $IC_{50}$ = 1,827 ± 337.24 nM<br>$K_i$ = 1,613 ± 60.97 nM |
| H172K | 3 | $K_m$ = 55.53 ± 2.57 µM<br>$V_{max}$ = 16.12 ± 0.29 nM/s<br>$k_{cat}$ = 0.0107 $S^{-1}$<br>$k_{cat}/K_m$ = 194 $S^{-1}M^{-1}$ | | | $IC_{50}$ = 1,030 ± 50.53 nM |
| H172R | 1 | $K_m$ = 1.09 ± 0.15 µM<br>$V_{max}$ = 1.19 ± 0.037 nM/s<br>$k_{cat}$ = 0.00036 $S^{-1}$<br>$k_{cat}/K_m$ = 262 $S^{-1}M^{-1}$ | | | $IC_{50}$ >10 µM |
| H172G | 1 | $K_m$ = 74.20 ± 2.31 µM<br>$V_{max}$ = 9.95 ± 0.14 nM/s<br>$k_{cat}$ = 0.0119 $S^{-1}$<br>$k_{cat}/K_m$ = 161 $S^{-1}M^{-1}$ | | | $IC_{50}$ = 810.2 ± 19.30 nM |
| H172C | 1 | $K_m$ = 66.79 ± 3.00 µM<br>$V_{max}$ = 12.59 ± 0.24 nM/s<br>$k_{cat}$ = 0.0094 $S^{-1}$<br>$k_{cat}/K_m$ = 141 $S^{-1}M^{-1}$ | | | $IC_{50}$ = 3,056 ± 158.3 nM |
| Q189K | 168 | $K_m$ = 41.45 ± 2.66 µM<br>$V_{max}$ = 39.78 ± 1.01 nM/s<br>$k_{cat}$ = 0.16 $S^{-1}$<br>$k_{cat}/K_m$ = 3,800 $S^{-1}M^{-1}$ | $IC_{50}$ = 73.65 ± 4.09 nM<br>$K_i$ = 64.09 ± 3.45 nM | $IC_{50}$ = 37.23 ± 0.83 nM<br>$K_i$ = 14.35 ± 1.65 nM | $IC_{50}$ = 39.31 ± 1.92 nM<br>$K_i$ = 14.60 ± 1.53 nM |
| Q189F | 38 | $K_m$ = 26.02 ± 2.40 µM<br>$V_{max}$ = 19.49 ± 1.16 nM/s<br>$k_{cat}$ = 0.3118 $S^{-1}$<br>$k_{cat}/K_m$ = 11,985 $S^{-1}M^{-1}$ | | | $IC_{50}$ = 34.02 ± 1.47 nM |
| Q189R | 27 | $K_m$ = 42.63 ± 2.72 µM<br>$V_{max}$ = 21.84 ± 0.52 nM/s<br>$k_{cat}$ = 0.1092 $S^{-1}$<br>$k_{cat}/K_m$ = 2,562 $S^{-1}M^{-1}$ | | | $IC_{50}$ = 37.82 ± 1.16 nM |
| Q189P | 18 | $K_m$ = 29.57 ± 2.25 µM<br>$V_{max}$ = 19.07 ± 0.97 nM/s<br>$k_{cat}$ = 0.3051 $S^{-1}$<br>$k_{cat}/K_m$ = 10,319 $S^{-1}M^{-1}$ | | | $IC_{50}$ = 14.08 ± 0.70 nM |
| Q189H | 20 | $K_m$ = 42.83 ± 3.24 µM<br>$V_{max}$ = 18.05 ± 0.51 nM/s<br>$k_{cat}$ = 0.1444 $S^{-1}$<br>$k_{cat}/K_m$ = 3,371 $S^{-1}M^{-1}$ | | | $IC_{50}$ = 32.71 ± 1.04 nM |
| Q189L | 17 | $K_m$ = 47.00 ± 3.83 µM<br>$V_{max}$ = 15.42 ± 0.48 nM/s<br>$k_{cat}$ = 0.0617 $S^{-1}$<br>$k_{cat}/K_m$ = 1,312 $S^{-1}M^{-1}$ | | | $IC_{50}$ = 39.25 ± 1.47 nM |

| | | | | | | |
|---|---|---|---|---|---|---|
| Q189S | 8 | $K_m$ = 27.97 ± 3.26 µM<br>$V_{max}$ = 17.28 ± 0.66 nM/s<br>$k_{cat}$ = 0.1728 $S^{-1}$<br>$k_{cat}/K_m$ = 6,178 $S^{-1}M^{-1}$ | | | $IC_{50}$ = 25.34 ± 1.72 nM | |
| Q189E | 8 | $K_m$ = 23.03 ± 1.71 µM<br>$V_{max}$ = 29.75 ± 1.91 nM/s<br>$k_{cat}$ = 0.4760 $S^{-1}$<br>$k_{cat}/K_m$ = 20,669 $S^{-1}M^{-1}$ | $IC_{50}$ = 80.19 ± 6.31 nM<br>$K_i$ = 36.86 ± 2.97 nM | $IC_{50}$ = 42.34 ± 3.07 nM<br>$K_i$ = 1.34 ± 0.14 nM | $IC_{50}$ = 49.71 ± 3.19 nM<br>$K_i$ = 4.51 ± 0.29 nM | |
| Q192T | 187 | $K_m$ = 32.94 ± 2.96 µM<br>$V_{max}$ = 19.94 ± 0.62 nM/s<br>$k_{cat}$ = 0.0399 $S^{-1}$<br>$k_{cat}/K_m$ = 1,200 $S^{-1}M^{-1}$ | $IC_{50}$ = 237.2 ± 17.12 nM<br>$K_i$ = 119.7 ± 6.57 nM | $IC_{50}$ = 185.5 ± 13.90 nM<br>$K_i$ = 71.40 ± 4.26 nM | $IC_{50}$ = 102.6 ± 5.52 nM<br>$K_i$ = 45.69 ± 4.43 nM | GC-376: 8DGB |
| Q192K | 62 | $K_m$ = 38.29 ± 2.05 µM<br>$V_{max}$ = 12.21 ± 0.23 nM/s<br>$k_{cat}$ = 0.0122 $S^{-1}$<br>$k_{cat}/K_m$ = 319 $S^{-1}M^{-1}$ | $IC_{50}$ = 303.6 ± 27.42 nM<br>$K_i$ = 430.3 ± 22.35 nM | $IC_{50}$ = 324.0 ± 18.93 nM<br>$K_i$ = 299.5 ± 23.79 nM | $IC_{50}$ = 219.1 ± 9.62 nM<br>$K_i$ = 88.90 ± 4.88 nM | |
| Q192S | 29 | $K_m$ = 46.24 ± 2.09 µM<br>$V_{max}$ = 28.69 ± 0.49 nM/s<br>$k_{cat}$ = 0.0574 $S^{-1}$<br>$k_{cat}/K_m$ = 1,241 $S^{-1}M^{-1}$ | $IC_{50}$ = 319.0 ± 12.02 nM<br>$K_i$ = 227.9 ± 17.06 nM | $IC_{50}$ = 183.9 ± 8.16 nM<br>$K_i$ = 82.32 ± 4.23 nM | $IC_{50}$ = 217.3 ± 10.59 nM<br>$K_i$ = 75.56 ± 8.45 nM | |
| Q192L | 12 | $K_m$ = 62.36 ± 3.76 µM<br>$V_{max}$ = 31.72 ± 0.79 nM/s<br>$k_{cat}$ = 0.1586 $S^{-1}$<br>$k_{cat}/K_m$ = 2,543 $S^{-1}M^{-1}$ | $K_i$ = 165.3 ± 17.86 nM | $K_i$ = 25.39 ± 1.81 nM | $IC_{50}$ = 121.9 ± 5.51 nM<br>$K_i$ = 70.99 ± 7.02 nM | |
| Q192R | 11 | $K_m$ = 59.39 ± 4.71 µM<br>$V_{max}$ = 17.70 ± 0.57 nM/s<br>$k_{cat}$ = 0.0283 $S^{-1}$<br>$k_{cat}/K_m$ = 477 $S^{-1}M^{-1}$ | | | $IC_{50}$ = 229.6 ± 11.54 nM | |
| Q192A | 9 | $K_m$ = 59.42 ± 3.02 µM<br>$V_{max}$ = 35.25 ± 0.0.73 nM/s<br>$k_{cat}$ = 0.1058 $S^{-1}$<br>$k_{cat}/K_m$ = 1,780 $S^{-1}M^{-1}$ | $K_i$ = 206.8 ± 14.64 nM | $K_i$ = 62.84 ± 5.63 nM | $IC_{50}$ = 140.7 ± 3.93 nM<br>$K_i$ = 58.10 ± 5.67 nM | |
| Q192I | 8 | $K_m$ = 52.72 ± 4.87 µM<br>$V_{max}$ = 21.47 ± 0.78 nM/s<br>$k_{cat}$ = 0.1031 $S^{-1}$<br>$k_{cat}/K_m$ = 1,955 $S^{-1}M^{-1}$ | $K_i$ = 179.8 ± 10.86 nM | $K_i$ = 41.16 ± 5.37 nM | $IC_{50}$ = 109.9 ± 2.45 nM<br>$K_i$ = 43.53 ± 3.13 nM | |
| Q192P | 8 | $K_m$ = 55.88 ± 3.74 µM<br>$V_{max}$ = 20.34 ± 0.55 nM/s<br>$k_{cat}$ = 0.0814 $S^{-1}$<br>$k_{cat}/K_m$ = 1,456 $S^{-1}M^{-1}$ | $K_i$ = 217.1 ± 19.24 nM | $K_i$ = 80.95 ± 8.68 nM | $IC_{50}$ = 135.1 ± 7.17 nM<br>$K_i$ = 76.19 ± 5.66 nM | |
| Q192N | 8 | $K_m$ = 54.93 ± 3.37 µM<br>$V_{max}$ = 9.19 ± 0.22 nM/s<br>$k_{cat}$ = 0.0408 $S^{-1}$<br>$k_{cat}/K_m$ = 744 $S^{-1}M^{-1}$ | | | $IC_{50}$ = 73.69 ± 4.96 nM | |
| Q192G | 7 | $K_m$ = 68.11 ± 2.55 µM<br>$V_{max}$ = 20.69 ± 0.33 nM/s<br>$k_{cat}$ = 0.0414 $S^{-1}$ | $K_i$ = 277.6 ± 19.21 nM | $K_i$ = 78.78 ± 7.01 nM | $IC_{50}$ = 319.1 ± 12.73 nM<br>$K_i$ = 106.3 ± 5.84 nM | |

| | | | | | |
|---|---|---|---|---|---|
| | | $k_{cat}/K_m$ = 608 $S^{-1}M^{-1}$ | | | |
| Q192H | 7 | $K_m$ = 60.51 ± 3.86 µM<br>$V_{max}$ = 27.12 ± 0.71 nM/s<br>$k_{cat}$ = 0.0814 $S^{-1}$<br>$k_{cat}/K_m$ = 1,345 $S^{-1}M^{-1}$ | $K_i$ = 237.2 ± 21.98 nM | $K_i$ = 61.19 ± 3.93 nM | $IC_{50}$ = 169.4 ± 7.27 nM<br>$K_i$ = 80.69 ± 7.33 nM |
| Q192Y | 7 | $K_m$ = 9.18 ± 0.62 µM<br>$V_{max}$ = 2.64 ± 0.12 nM/s<br>$k_{cat}$ = 0.0004 $S^{-1}$<br>$k_{cat}/K_m$ = 43 $S^{-1}M^{-1}$ | | | $IC_{50}$ >10 µM |
| Q192V | 6 | $K_m$ = 59.77 ± 3.01 µM<br>$V_{max}$ = 18.36 ± 0.38 nM/s<br>$k_{cat}$ = 0.0734 $S^{-1}$<br>$k_{cat}/K_m$ = 1,229 $S^{-1}M^{-1}$ | $IC_{50}$ = 136.1 ± 5.66 nM<br>$K_i$ = 122.0 ± 9.62 nM | $IC_{50}$ = 82.17 ± 2.54 nM<br>$K_i$ = 78.46 ± 6.47 nM | $IC_{50}$ = 95.58 ± 4.23 nM<br>$K_i$ = 30.87 ± 1.12 nM |
| Q192W | 5 | $K_m$ = 50.15 ± 4.32 µM<br>$V_{max}$ = 24.00 ± 0.80 nM/s<br>$k_{cat}$ = 0.0691 $S^{-1}$<br>$k_{cat}/K_m$ = 1,378 $S^{-1}M^{-1}$ | $K_i$ = 149.0 ± 12.78 nM | $K_i$ = 32.88 ± 2.89 nM | $IC_{50}$ = 65.94 ± 2.54 nM<br>$K_i$ = 43.65 ± 2.32 nM |
| Q192E | 5 | $K_m$ = 60.37 ± 3.88 µM<br>$V_{max}$ = 32.98 ± 0.87 nM/s<br>$k_{cat}$ = 0.0660 $S^{-1}$<br>$k_{cat}/K_m$ = 1,093 $S^{-1}M^{-1}$ | $K_i$ = 296.1 ± 21.17 nM | $K_i$ = 39.21 ± 2.92 nM | $IC_{50}$ = 193.1 ± 9.27 nM<br>$K_i$ = 114.4 ± 10.82 nM |
| Q192C | 3 | $K_m$ = 55.53 ± 3.46 µM<br>$V_{max}$ = 21.80 ± 0.54 nM/s<br>$k_{cat}$ = 0.0872 $S^{-1}$<br>$k_{cat}/K_m$ = 1,570 $S^{-1}M^{-1}$ | $K_i$ = 147.7 ± 16.25 nM | $K_i$ = 27.98 ± 1.58 nM | $IC_{50}$ = 96.82 ± 2.40 nM<br>$K_i$ = 54.12 ± 2.79 nM |
| Q192D | 3 | $K_m$ = 68.35 ± 5.76 µM<br>$V_{max}$ = 18.58 ± 0.67 nM/s<br>$k_{cat}$ = 0.0372 $S^{-1}$<br>$k_{cat}/K_m$ = 544 $S^{-1}M^{-1}$ | | | $IC_{50}$ = 271.5 ± 17.63 nM |
| Q192F | 2 | $K_m$ = 54.41 ± 4.34 µM<br>$V_{max}$ = 42.44 ± 1.35 nM/s<br>$k_{cat}$ = 0.1698 $S^{-1}$<br>$k_{cat}/K_m$ = 3,120 $S^{-1}M^{-1}$ | $K_i$ = 137.5 ± 11.09 nM | $K_i$ = 21.41 ± 2.28 nM | $IC_{50}$ = 92.68 ± 6.53 nM<br>$K_i$ = 85.50 ± 12.51 nM |
| Q189E/ H172Y | 0 | $K_m$ = 61.34 ± 4.83 µM<br>$V_{max}$ = 24.76 ± 0.80 nM/s<br>$k_{cat}$ = 0.0619 $S^{-1}$<br>$k_{cat}/K_m$ = 1,009 $S^{-1}M^{-1}$ | $IC_{50}$ = 652.7 ± 30.98 nM<br>$K_i$ = 618.2 ± 36.42 nM | $IC_{50}$ = 169.9 ± 11.54 nM<br>$K_i$ = 83.84 ± 4.61 nM | $IC_{50}$ = 603.7 ± 17.42 nM<br>$K_i$ = 528.5 ± 47.25 nM |

[a]The occurrence of mutation was analyzed using the GISAID CoVsurver as of July 2nd, 2022. $k_{cat}$, $V_{max}$, $K_m$, $k_{cat}/K_m$, $IC_{50}$ and $K_i$ values are the average of two repeats. Enzymatically inactive mutants are colored in yellow. $M^{pro}$ mutants that have comparable enzymatic activity as the WT ($k_{cat}/K_m$ <10-fold change) and are resistant to nirmatrelvir ($K_i$ > 10-fold increase) are colored in red.

### Table S2. X-ray Data Collection and Refinement Statistics

| **Data Collection** | | | | | | | | | | | |
|---|---|---|---|---|---|---|---|---|---|---|---|
| Structure (PDB ID) | 8D4J | 8D4K | 8D4L | 8D4M | 8D4N | 8DFN | 8DD1 | 8DFE | 8DD9 | 8DGB | 8DCZ |
| Mutation | H172Y | H172Y | S144A | S144A | E166Q | H164N | H164N | S144L | S144L | Q192T | M165Y |
| Ligand | Apo | GC376 | Apo | GC376 | Apo | Apo | GC376 | Apo | GC376 | GC376 | Nirmatrelvir |
| Space Group | $P2_1$ | C2 | $P2_1$ | C2 | $P2_1$ | $P2_1$ | C2 | C2 | C2 | P1 | $P2_1$ |
| Cell Dimensions | | | | | | | | | | | |
| *a*, *b*, *c* (Å) | 44.962 | 114.549 | 44.625 | 114.134 | 44.816 | 44.668 | 113.899 | 113.554 | 113.707 | 47.03 | 45.566 |
| | 53.474 | 53.111 | 53.707 | 53.653 | 53.687 | 53.738 | 53.55 | 54.075 | 53.153 | 53.241 | 53.786 |
| | 114.185 | 45.643 | 114.685 | 45.457 | 114.532 | 114.781 | 45.321 | 44.615 | 45.345 | 59.532 | 114.76 |
| a, b, g (°) | 90 | 90 | 90 | 90 | 90 | 90 | 90 | 90 | 90 | 67 | 90 |
| | 101.14 | 102.67 | 101.3 | 101.97 | 101.79 | 100.81 | 101.9 | 100.73 | 102.22 | 79.57 | 100.53 |
| | 90 | 90 | 90 | 90 | 90 | 90 | 90 | 90 | 90 | 88.8 | 90 |
| Resolution (Å) | 50-1.78 | 50-1.89 | 50-1.70 | 50-1.81 | 50-2.70 | 50-2.04 | 50-2.04 | 50-1.89 | 50-2.04 | 50-2.87 | 50-2.38 |
| No. Reflections | 51114 | 21112 | 56762 | 24143 | 14631 | 33845 | 16735 | 20077 | 16416 | 10878 | 21947 |
| $R_{merge}$ (%) | 6.3 | 8.4 | 5.5 | 7.4 | 17.2 | 8.7 | 7.8 | 7.1 | 9.7 | 7.9 | 9.2 |
| *I* / *óI* | 24.2 (2.73) | 20.35 (2.01) | 28.72 (3.46) | 21.77 (2.34) | 9.13 (2.02) | 16.75 (2.14) | 22.18 (2.27) | 26.7 (2.63) | 13.59 (2.31) | 9.03 (3.06) | 12.33 (2.13) |
| Completeness (%) | 99.5 | 98.6 | 96.8 | 98.1 | 99.1 | 98.6 | 97.0 | 94.0 | 96.9 | 93.5 | 98.6 |
| Redundancy | 3.5 (2.6) | 4.1 (3.9) | 3.4 (3.2) | 3.9 (4.1) | 3.7 (3.6) | 5.2 (5.2) | 4.9 (4.7) | 6.2 (5.9) | 3.7 (3.3) | 2.2 (1.5) | 4.0 (3.5) |
| **Refinement** | | | | | | | | | | | |
| Resolution (Å) | 38.71-1.78 | 48.02-1.89 | 38.87-1.70 | 48.41-1.81 | 38.80-2.70 | 43.79-2.04 | 34.24-2.04 | 48.41-1.89 | 39.92-2.04 | 48.98-2.87 | 39.28-2.38 |
| $R_{work}/R_{free}$ (%) | 16.6/20.9 | 16.6/22.4 | 17.1/21.3 | 17.2/22.9 | 20.6/25.8 | 19.1/25.1 | 18.5/24.8 | 20.8/26.8 | 16.5/22.8 | 22.7/27.2 | 19.2/25.1 |
| **No. Heavy Atoms** | | | | | | | | | | | |
| Protein | 4912 | 2443 | 4912 | 2431 | 4844 | 4703 | 2345 | 2421 | 2411 | 4723 | 4757 |
| Ligand/Ion | 6 | 29 | 0 | 29 | 0 | 0 | 29 | 0 | 29 | 58 | 70 |
| Water | 478 | 187 | 471 | 246 | 40 | 304 | 148 | 201 | 208 | 11 | 140 |
| ***B*-Factors (Å$^2$)** | | | | | | | | | | | |
| Protein | 27.19 | 30.87 | 25.91 | 31.04 | 34.42 | 33.18 | 45.22 | 30.96 | 33.51 | 56.10 | 40.08 |
| Ligand/Ion | 36.91 | 30.63 | 0 | 32.29 | 0 | 0 | 40.87 | 0 | 25.19 | 59.20 | 31.48 |
| Water | 34.53 | 37.51 | 33.21 | 37.49 | 20.3 | 36.22 | 48.83 | 34.38 | 38.29 | 54.78 | 37.38 |
| **Ramanchandran Plot** | | | | | | | | | | | |
| Favored Region (%) | 98.5 | 98.0 | 98.5 | 98.7 | 98.7 | 98.3 | 98.3 | 98.7 | 98.7 | 97.4 | 97.7 |
| Allowed Region (%) | 1.5 | 2.0 | 1.5 | 1.3 | 1.0 | 1.3 | 1.3 | 1.3 | 1.3 | 2.6 | 2.0 |
| Outlier Region (%) | 0.0 | 0.0 | 0.0 | 0.0 | 0.3 | 0.3 | 0.3 | 0.0 | 0.0 | 0.0 | 0.3 |

*values in the parentheses represent highest resolution shells
